# Mobile genetic elements are active and responsive to community context in model microbial consortium

**DOI:** 10.64898/2026.09.08.750204

**Authors:** Lillian C. Lowrey, Josue A Rodriguez-Ramos, Amy E. Zimmerman, William C. Nelson, James D. Jaryenneh, Kelly P. Williams, Kirsten S. Hofmockel, Joseph Schoeniger, Catherine M. Mageeney

## Abstract

Insertion and excision of genomic islands (GIs), chromosomally-integrated mobile genetic elements (MGEs), are major sources of microbial genome plasticity and can impact gene expression and phenotype of the host organism. GI mobilization also influences microbial communities beyond the host organism as GI excision generates MGEs that can be transferred between community members through horizontal gene transfer and induction of prophages can kill host populations, which impacts community structure. Established computational methods now enable precise GI mapping in genomes, as well as highly sensitive detection of GI excision from deep-genome sequencing data. We applied these approaches to metagenomic datasets from a defined soil microbial consortium grown on glass beads under hydration stress and compared GI activity with that observed in monoculture. Under these environmentally structured community growth conditions, GI excision was more abundant and involved a broader range of host species and GI types than under isolate growth conditions. Combined analysis with metatranscriptomic and metaproteomic data identified patterns of GI gene expression associated with induction. Three GIs showed particularly high excision together with strong transcription, numerous detected proteins, and evidence of association with potential transfer particles, including phages or vesicles. These results indicate that isolate studies can miss a substantial environmentally responsive layer of GI activity. More broadly, this work establishes a framework for mining existing community multi-omic datasets to quantify dynamic genome restructuring, identify active but poorly understood GIs, and generate mechanistic hypotheses about the processes that shape microbial genome plasticity and gene flow.

**Significance statement:** Genomic island (GI) mobilization is a major source of microbial genome plasticity, yet has mostly been studied in isolates, leaving GI behavior in environmentally structured communities poorly understood. In a model soil consortium, we show that environmentally-relevant conditions elicit substantially more abundant and broad GI excision than isolate cultures, indicating that conventional studies can miss an important layer of microbial genome dynamics. By linking excision to transcription, protein production, and candidate transfer particles, this approach opens a route to studying active GIs whose mobilization mechanisms are unknown. Mining existing environmental multi-omic datasets in this way could improve ecosystem models and inform safer, more predictable microbial engineering and improved biocontainment.

## INTRODUCTION

In nature, microbial communities are constantly challenged by environmental fluctuations, including hydration, organic and inorganic nutrients, oxygen, physical proximity to other microbes, and numerous other stressors. Mobile genetic elements (MGEs) represent an opportunity for coping with environments under flux more rapidly than random genetic drift. Through horizontal gene transfer (HGT), MGEs can bestow new metabolic or defensive capabilities (e.g. antibiotic resistance, increased pathogenicity, bolstered stress tolerance) to the recipient organism (1). Thus, MGE mobilization shapes both microbial ecology and evolution by impacting the functional potential, diversity, and stability of whole communities (2). MGE mobilization is tightly regulated because it carries risks associated with loss of genetic information from the chromosome, disruption of coding or regulatory sequences, or changes in gene dosage that can impact fitness during excision or integration. However, the cues that activate MGE excision are only well understood for a few model isolates, which point to stress response and nutrient limitation as key factors (1, 3)).(1, 3)

Many MGEs carry site-specific recombinases (i.e. integrases) and the cognate DNA sequences (attachment (*att*) sites) that enable their integration into the chromosome as genomic islands (GIs). *Att* sites provide a tractable way to directly measure GI mobilization within community datasets. GIs integrate at defined attachment sites within bacterial chromosomes (*attL* and *attR*), while GI excision creates detectable junctions on the bacterial chromosome (*attB*) and the GI (*attP*) (4). Recently developed bioinformatic tools take advantage of these features to quantify signatures of GI mobilization using (meta)genomic sequencing alone, enabling precise GI prediction, categorization, and induction tracking (5–9). Coupling metagenomic data analysis with evidence of MGE expression from corresponding RNA and protein data can provide deeper insights into the activation of each GI and aid prediction of transfer mechanisms. Harnessing new computational tools to track genomic island dynamics within non-model microbial communities while accounting for environmental context allows for the precise characterization of site-specific integration events, driving breakthroughs in synthetic biology platforms across gene therapy, advanced biologics, and engineered cell factories (10–14).

In this work, we applied a unique computational toolset to precisely predict and track GI excision, representative of GI mobilization, under environmentally relevant stress conditions by combining genomic, metagenomic, metatranscriptomic, and metaproteomic datasets from a defined microbial community. We comprehensively predicted GIs and transposons in an established 39-member Model Soil Consortium (MSC-1), which includes eight taxonomically diverse isolates that form a naturally co-occurring sub-community capable of chitin decomposition (15, 16). We then quantified and compared GI excision events for the nested isolates under conventional mitomycin C induction in isolation and environmentally relevant community growth conditions. We show that community context greatly increases GI activation and produces a more accurate estimation of GI mobilization in the natural soil environment than conventional analyses. Further analysis revealed multiple active prophages that produce visually confirmed phage particles and a GI showing evidence of vesicle-mediated transfer. This work underscores the importance of community context in MGE ecology and demonstrates a computational approach for moving from static MGE prediction to quantification of MGE dynamics within non-model and wild microbial communities (**Figure 1**).

**Figure 1.**
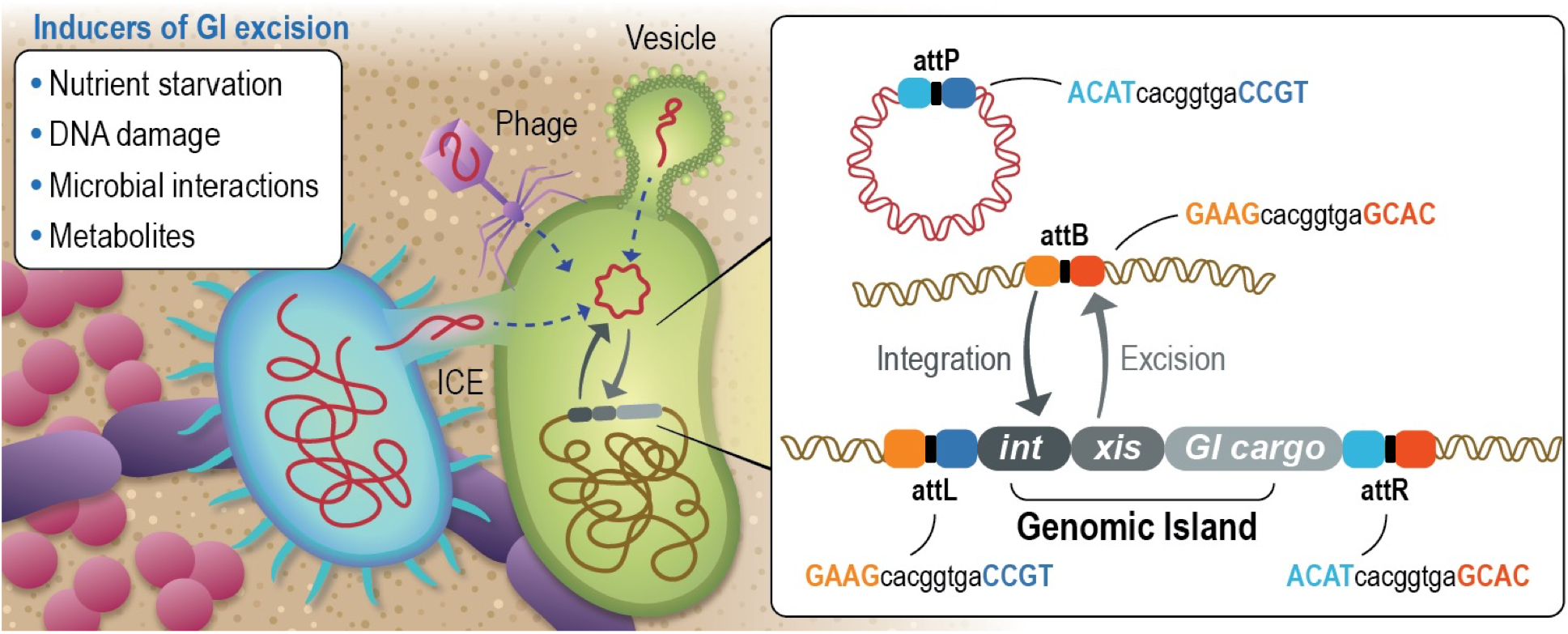
Genomic islands, regions of DNA that integrate and excise from genomes, are horizontally transferred between members of a microbial community and can be tracked using multi-omics datasets. Mobile genetic elements (MGEs), such as plasmids and integrative conjugative elements (ICEs) are transferred horizontally within bacterial communities by phage, vesicles, or conjugation with neighboring cells (left panel). Integrase-mediated recombination between genomic *attB* sites and MGE *attP* sites integrates some MGEs into the genome as genomic islands (GIs), with *attL* and *attR* sites formed at the GI boundaries (right panel). GIs excise using an integrase and excisionase protein in response to a variety of signals, such as nutrient starvation, DNA damage, microbial interactions, and metabolites, reproducing the *attP* and *attB* sequences. Because the boundaries of each *att* site differ, the occurrence of each within genome or metagenome sequencing reads was counted to track GI excision in this work. Typically, in addition to carrying the integrase- and excisionase-encoding genes (*int* and *xis* respectively), GIs also carry a variety of cargo that may impact cell phenotype, which was investigated here using metatranscriptomic and metaproteomic datasets.

## RESULTS

### DNA recombination is evident in the MSC-1 isolates and elevated under community growth conditions

We analyzed (meta)genomic data from both monocultures of the eight MSC-1 isolates that were treated with mitomycin-C (MMC) and incubations of the complete MSC-1 community with different levels of hydration in an inert porous matrix intended to mimic soil structure (17) to compare rates of mobilization by transposition, homologous recombination, and non-homologous end joining (18, 19). Recombined sequences were identified by mapping whole genome sequencing reads to the reference genomes and sorted into 500 base pair (bp) bins to create histograms of recombination hotspots across the genomes (**Figure 2** and **Supplemental Figure 1**). At 24 hours post MMC treatment, the *Sphingopyxis* genome underwent a high rate of recombination (**Figure 2D**) with 3551 recombination events per million sequencing reads, 7x higher than the average for the other isolates (437 events per million sequencing reads) (**Supplemental Table 1**). We observed that the amount of DNA recombination was elevated across all isolates during growth in community compared to monoculture (**Figure 2** and **Supplemental Figure 1**).

**Figure 2.**
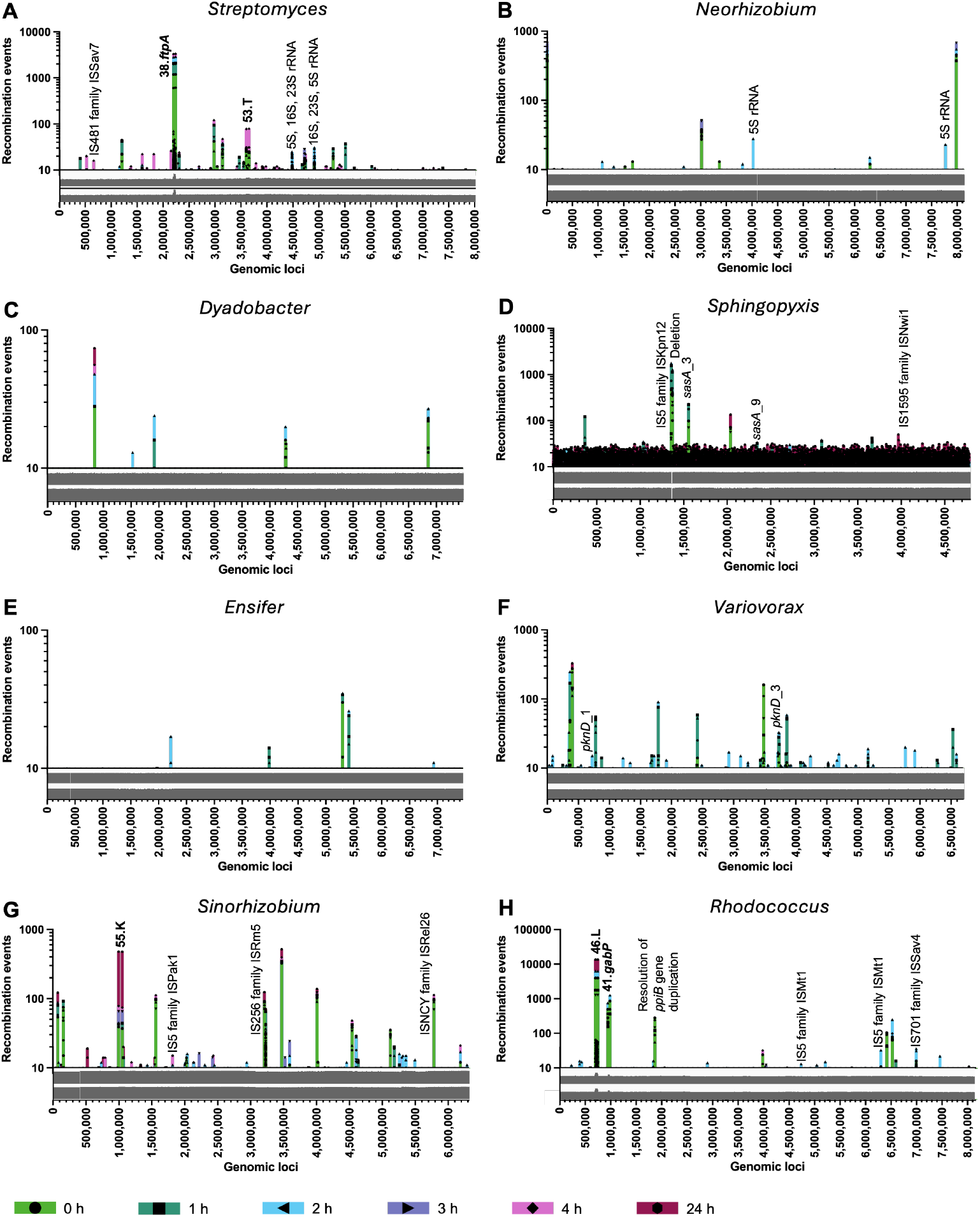
DNA damage leads to recombination at homologous sequences, IS-elements, and GIs in MSC-1 isolates grown in monoculture. DNA recombination events were identified by high-depth, short-read sequencing, sorted into 500 bp bins spanning the genome of (**A**) *Streptomyces*, (**B**) *Neorhizobium*, (**C**) *Dyadobacter*, (**D**) *Sphingopyxis*, (**E**) *Ensifer*, (**F**) *Variovorax*, (**G**) *Sinorhizobium*, and (**H**) *Rhodococcus* isolates grown in monocultures induced with MMC. IS-element and GI recombination events are labeled with the IS-elements name and family or GI name, respectively. Suspected homologous recombination events are labeled with the homologous genes. Multiple time points following MMC treatment to induce stress from DNA damage are shown (0, 1, 2, 3, 4, and 24 hours). Log-scale graphs of DNA sequencing coverage aligned to the forward (top) or reverse (bottom) strand are shown below each plot. The range for each plot spans 0 – 1,000 reads for all organisms other than *Streptomyces* and *Rhodococcus*, which have maximums of 10,000 reads.

A high level of recombination was observed in regions with identified GIs (53.T, 38.ftpA, and 5.fumA in *Streptomyces*, 55.K in *Sinorhizobium*, and 48.L and 41.*gabP* in *Rhodococcus*) (**Figure 2A, G, and H** and **Supplemental Figure 1 A and H**). Other DNA recombination events, unrelated to identified GIs, were also revealed, including putative non-allelic homologous recombination (HR) as indicated by pairs of recombination peaks in homologous sequences. Recombination within rRNA operons, known hotspots for homologous recombination, was observed in the *Streptomyces, Rhodococcus*, and *Neorhizobium* isolates in monoculture (**Figure 2A and B**) and in all 8 isolates when incubated as the complete MSC-1 community (**Supplemental Figure 1**) (20–24).

The recombination histograms also revealed a variety of mobilizing insertion sequence (IS) elements with most belonging to families known to conduct primarily replicative or copy-and-paste transposition (25) (**Figure 2 and Supplemental Figure 1**). Therefore, some spurious peaks in the histograms could be created by insertion of a copy of these IS-elements into new genomic loci. IS-elements may also provide DNA substrates for homologous recombination (26), as was likely the case for a pair of directly oriented ISMt1 elements in *Rhodococcus* (**Figure 2H**).

### Computationally predicted GIs are abundant and distinct within MSC-1

TIGER and Islander (5, 27, 28) were used to precisely predicted 121 GIs in the MSC-1 genomes (**Supplemental Tables 2 and 3**). The MSC-1 isolates contained 40 GIs predicted as 9 prophages, 6 defective prophages, 3 ICEs, 7 GIs with metabolism genes, and 2 defense islands (**Supplemental Table 2, Figure 3A**). Insufficient annotation and unresolved mechanisms of mobility prevented typing of the remaining 13 GIs according to gene content. We additionally predicted 37 transposons within the genomes of the MSC-1 isolates using TIGER (**Supplemental Table 4**).

**Figure 3.**
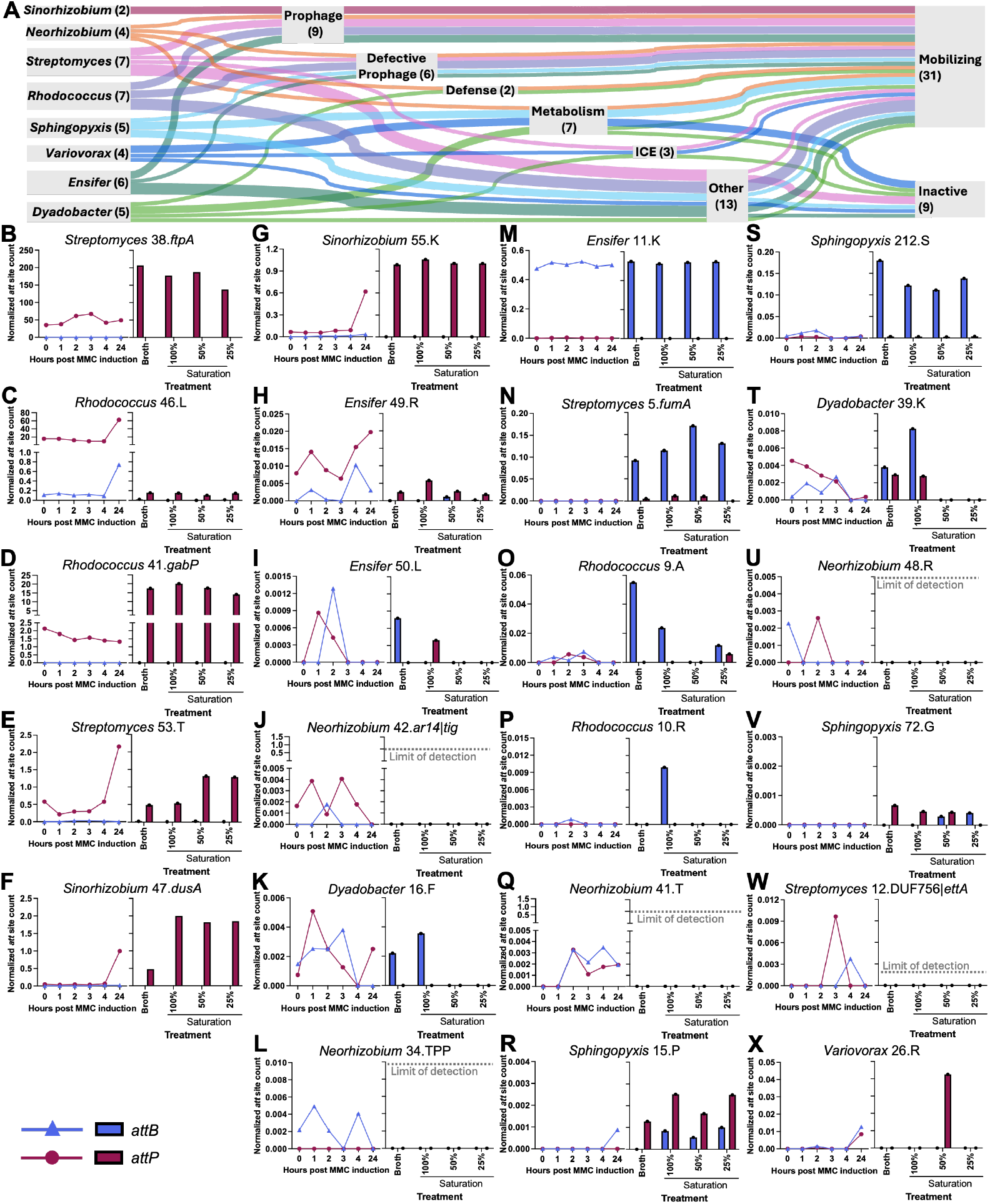
Precise GI mobilization within MSC-1 isolates. (**A**) Sankey diagram of the bacterial source, type, and activity of the identified genomic islands in MSC-1 isolates. The number of GIs in each category is shown in parentheses. (**B-X**) Normalized *attP* (red circles/bars) and *attB* (blue triangles/bars) counts for the mobilizing GIs in monoculture following MMC treatment (left, line graph) and community growth with a porous matrix and variable hydration levels (right, grouped bar graph) for (**B-J**) prophage islands, (**K, L**) defense islands, (**M-R**) defective prophage islands, (**S-V**) metabolism islands, and (**W, X**) ICEs. For each GI, Y-axis scales and units are identical between the paired graphs. Key at bottom left. Coculture growth in liquid medium is indicated as “Broth”.

Most predicted GIs from MSC-1 isolates (33/40) had unique gene content (as determined by AAI clustering analysis), allowing accurate attribution of reads to these GIs and the observation that all excision events could be mapped to a single GI within MSC-1. Four GIs found in MSC-1 isolates shared similarity with GIs in other consortium members (**Supplemental Figure 2**). However, no significant sequence similarity was observed at the nucleotide level except for two GIs in *Rhodococcus* (2.Hyp and 3.Hyp). These share the same *att* site therefore excision events cannot be accurately attributed (**Supplemental Figure 3**).

### Prophage, defective prophage, defense, metabolic, and ICE GIs excise from the MSC-1 isolates in response to DNA damage and hydration stress

To sensitively quantify GI excision, the number of whole genome sequencing reads containing the predicted attachment sites (*attB, attP, attL*, and *attR*) was counted for each GI and normalized at a per genome level. We observed 31 GIs excise from the MSC-1 isolates (**Figure 3A**). All prophage, defective prophage, and defense islands excised (**Figure 3A**), while metabolism, ICE, and “other” islands excised less frequently (4/7, 2/3, and 8/13 respectively). As expected for prophage, GIs were excising and replicating (greater ratio of *attP* to *attB*), while defective prophage GIs were excising without replicating (greater ratio of *attB* than *attP*), likely due to GI decay hindering GI replication mechanisms.

Some of these functional and defective prophage GIs excised abundantly (greater ratio of *attP* than *attL* and *attR*), including 38.ftpA, 46.L, 41.gabP, 53.T, 47.dusA, 55.K, and 11.K, while others were less active, including 49.R, 50.L, 42.ar14|tis, 5.fumA, 9.A, 10.R, 41.T and 15.P, which were only present as circular molecules in an average of 1/25 cells. Because community members are naturally present in different proportions, some rare mobilization events were likely missed due to the depth of metagenome sequencing (**Figure 3 J, L, Q, and U**) (17).

The remaining mobilizing GIs carry defense genes (**Figure 3K, 2L**), metabolic genes (**Figure 3S-V**), are ICEs (**Figure 3W, 2X**), or cannot be typed (Other) (**Supplemental Figure 4**). While excision of these islands was less abundant, their mobilization likely impacts foreign DNA defense mechanisms or reshapes cell metabolism; and their uneven mobilization produces subpopulations with different genotypes and likely variable phenotypes.

We sought to understand what factors may contribute to GIs excision by comparing standard GI induction techniques (DNA damage-stressed monocultures) to more environmentally relevant stress conditions (cocultures grown under variable hydration levels to mimic moisture gradients found in natural soil systems). Primarily, functional and defective prophage GIs were excised by MMC DNA-damage stress (38.ftpA, 16.F, 41.T, 9.A, 12.DUF756|*ettA*, 7.L, 49.R, 46.L, 53.T, 47.dusA, 55.K, and 26.R), while excision was broader and typically more abundant under hydration stress (**Figure 3B-X**).

Community growth in a structured, porous matrix (analogous to soil) had a strong influence on GI excision compared to planktonic growth. 47.dusA, 49.R, 16.F, and 39.K excise more abundantly when MSC-1 was grown in a matrix (liquid medium with glass beads), while 38.ftpA 9.A, 212.S, 72.G, 6.dhmA, and 5.HYP|Y-Int excise most abundantly in growth in liquid medium only. Hydration levels showed mixed impacts on GI excision: 53.T exhibited a high level of excision and replication in drier conditions, while most moisture-responsive GIs (41.gabP, 49.R, 16.F, 9.A, 39.K, 6.dhmA, and 5.HYP|Y-Int) excised more abundantly at higher hydration levels, and 5.fumA, 26.R, 3.HYP|HYP, and 4.DUF1016 excised most abundantly at 50% saturation (**Figure 3 and Supplemental Figure 4**). These results could indicate different cellular responses (or taxon-specific adaptations) to moisture levels within the soil consortium. Taken together, these results broadly indicate that GIs respond differently to variable stress and growth conditions and evince the need to use environmentally relevant stress conditions when analyzing GI biology.

Due to similarities in medium and treatment, the impact of community context on GI excision was investigated by comparing initial (T0) monoculture to MSC-1 community growth in liquid medium (**Figure 3B-X**, T0 and liquid medium, respectively). We observed similar levels of excision of GIs 53.T, 11.K, 39.K, and 7.L regardless of community context. In contrast, cultivation in community increased excision of 38.*ftpA*, 41.*gabP*, 47.*dusA*, 9.A, 212.S, and 5.HYP|Y-Int. Strikingly, excision of 5.*fumA*,15.P, 72.G, and 6.dhmA was only observed in community culture. Prophage GIs 46.L, and to a lesser extent, 49.R, were less mobile under community growth. These results support the value of analyzing communities rather than isolates to accurately investigate environmentally relevant GI mobilization.

### Multi-omics validates GI excision and provides a deeper understanding of gene expression in response to hydration

Gene expression and protein production associated with GI activation was evaluated to understand the potential impacts of active GIs their host. We considered transcription at both the whole GI and individual gene levels (**Figure 4** and **Supplemental Figures 7-12**). Transcript levels were low from GIs found in *Neorhizobium, Sinorhizobium*, and *Variovorax* in all conditions, likely because these species are less abundant within the consortium and thus had lower read counts compared to the other MSC-1 isolates (17). Conversely, all the GIs in *Sphingopyxis* and *Ensifer* were highly expressed in all conditions.

**Figure 4.**
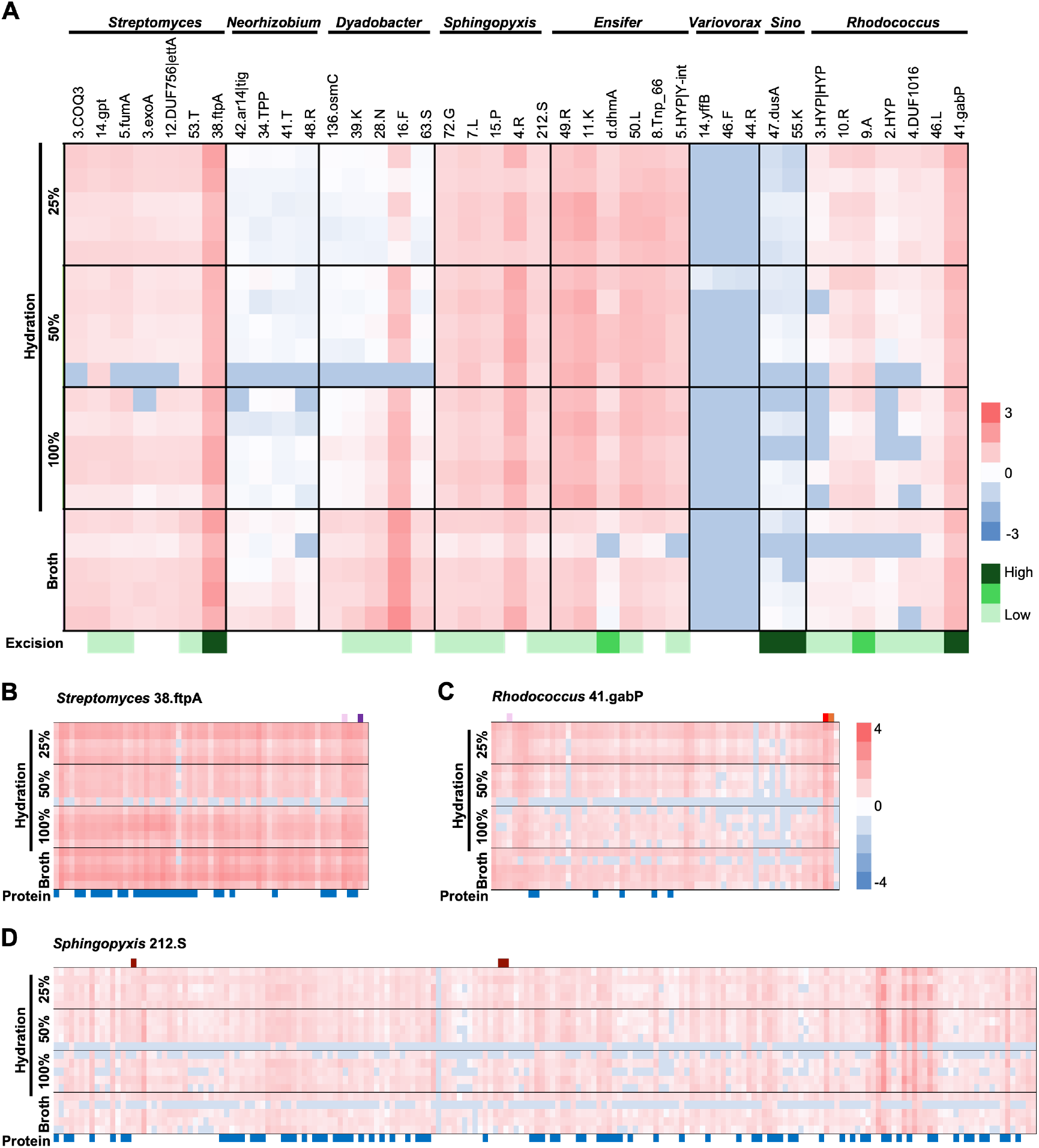
Global GI and gene-by-gene RNA and protein expression during MSC-1 community growth. **(A)** Global GI transcription during MSC-1 community growth. RNA expression (gene length corrected Trimmed Mean of M-values, geTMM) was summed across individual GIs for each taxon, then Z-score normalized across all GIs and conditions for visualization. The z-score range legend (red to blue) is on the right. Green boxes below report the rate of excision from **Figure 3B-X**. High is greater than 1, medium is 0.5-1, low is less than 0.5 events per genome. *Sinorhizobium* is abbreviated as *Sino*. **(B-D)** Gene-specific RNA expression and protein production from select GIs, including **(B)** *Streptomyces* 38.ftpA, **(C)** *Rhodococcus* 41.gabP, and **(D)** *Sphingopyxis* 212.S. RNA transcription is shown as Z-score normalized transcripts per million (TPM) for each gene predicted in each GI. Expression scale is on the right. Proteins of interest are indicated above the heatmap: purple = tyrosine integrase, orange = serine integrase, light orange = serine core integrase domains, maroon = transposase, pink = excisionase, red = repressor. Detection of encoded protein products by metaproteomics is shown in blue below the heatmap. Samples grown in liquid medium are referred to as ““Broth”.

High expression in *Sphingopyxis* corresponded to active excision for all GIs - except 4.R (**Figure 3 and Supplemental Figure 4**).

For the remaining MSC-1 isolates, global GI expression differed across the hydration levels in a GI-dependent manner. For example, *Streptomyces* 38.ftpA was transcriptionally active in all conditions, mirroring the high level of excision and replication observed of this GI (**Figure 2A** and **Figure 4A**). The remaining *Streptomyces* GIs have lower expression levels across all conditions. *Dyadobacter* GIs are transcribed highly in liquid media and moderately when grown in a porous matrix at 100% hydration, but expression decreases in 50% and 25% hydration suggesting GI expression positively correlates with hydration (**Figure 4A**). *Rhodococcus* 41.gabP is one of the most abundant GIs in the entire dataset confirmed by evidence of GI excision (**Figure 4A**).

We observed weak transcription of the predicted transposons, indicating minimal activity (**Supplemental Figure 5**). Exceptions included all five transposons predicted in *Ensifer* and *Rhodococcus* 10.Hyp|16S_ribosomal_RNA, which are highly expressed in all conditions. The *Streptomyces* and remaining *Rhodococcus* transposons are upregulated only when grown at lower moisture (25% hydration).

Metaproteomic data were evaluated to understand which GI-encoded proteins are produced during community growth. We note that while detection confirms the presence of the protein, absence does not exclude the possibility of that protein being translated. Frequently, proteins are produced below detection limits; this is routinely the case for non-structural proteins of phages (29–31). The same two prophages observed to have high rates of excision, 38.ftpA and 41.gabP, were also transcriptionally upregulated with multiple proteins evident. We observed 30 proteins from *Streptomyces* 38.ftpA, including the endolysin, exonucleases, transcriptional regulators, and many structural proteins (**Figure 4B**). *Rhodococcus* 41.gabP also has high levels of GI excision and replication and high levels of transcript across all genes with only two hypothetical genes downregulated, and multiple structural proteins observed (**Figure 4C**). MMC-induced filtrates from *Streptomyces* and *Rhodococcus* isolates contained Siphoviridae virions (**Supplemental Figure 6A and 6B**), confirming the active prophages can mediate transfer through viral particles.

*Sphingopyxis* 212.S, a metabolic GI, was also highly expressed across all conditions, with many detected proteins (**Figure 4D**). A majority of the proteins present on this GI are involved with cell wall synthesis, but it also carries operons for mannose and other sugar dehydrogenases with potential roles in energy generation and extracellular polysaccharide synthesis, as well as numerous metalloproteases which may be relevant for nutrient cycling (32). We observed 49 ± 6.9nm spherical structures (**Supplemental Figure 6C**) in filtrates from MMC-induced *Sphingopyxis*. We hypothesize these spherical structures are extracellular membrane vesicles that transport 212.S (and possible other GIs predicted in *Sphingopyxis*) to neighboring cells within the community, in line with transport mechanisms observed for several marine bacteria (33–36).

The remaining *Streptomyces* GIs had similar transcriptional profiles across the genes in the GI in liquid medium, 100% and 50% hydration conditions (**Supplemental Figure 7**). Conversely, at 25% hydration, positive z-scores indicated increased transcription of all genes across all GIs.

*Sphingopyxis* transcriptional activity was increased across all other predicted GIs, and there does not appear to be treatment-specific expression (**Supplemental Figure 8**). The remaining *Rhodococcus* GIs were expressed below the mean in liquid media, 100%, and 50% hydration conditions but weakly upregulated at 25% hydration, suggesting desiccation may activate these GIs (**Supplemental Figure 9**). We observed two proteins from these GIs, corresponding to the major capsid protein and the major tail protein of prophage 46.L.

*Ensifer* gene expression was higher than the mean for most genes across all treatments, and several proteins were observed, including the only excisionase protein in any MSC-1 isolate (49.R) (**Supplemental Figure 10**). Conversely, *Dyadobacter* GIs were only expressed above the mean in liquid medium (**Supplemental Figure 11**) suggesting desiccation may have a negative impact on GIs in *Dyadobacter*. The gene-by-gene analysis for *Neorhizobium, Sinorhizobium*, and *Variovorax* GIs was sparse, likely due to low read counts (**Supplemental Figure 12**). All GIs with attributable transcripts were expressed below the mean of the dataset and no proteins from these GIs were evident.

Overall, high levels of GI excision were supported by high levels of transcription and corresponding detection of proteins across that GI. We observed no integrases in the proteomics dataset; however, we see increased transcription of both the integrase and predicted excisionases across most GIs which corresponds to observations supporting excision from our metagenomic data.

## DISCUSSION

Here we present a powerful approach for identification and characterization of GI mobility within a microbial community. Using our precise GI prediction workflows (5, 27, 28, 37) coupled with multi-omics data, we were able to precisely and confidently predict 40 GIs in eight isolates from a model soil consortium (MSC-1) and show that most of these GIs (77%) are capable of excision. GI excision was more abundant and widespread when microbes were grown as a full community, with many GIs only excising when grown in the community context (7/31 GIs), suggesting interactions with other organisms likely induce GI activity. This work provides evidence to support the long-held hypothesis that HGT is more abundant in microbial communities than in isolates, likely due to increased cell-to-cell contact, increased presence of stressors from competitive interactions (i.e. nutrient limitation and production of secondary metabolites, etc.), and/or the stabilization of beneficial traits through gene drives (4, 38). Incorporation of multi-omics data increases confidence in both the occurrence of excision and the biological interpretation of GI “activation” (e.g., prophage particle formation versus non-lytic excision) (**Figure 1**). These methods could be applied to understand HGT within wild microbial communities to increase the potential for co-opting these diverse gene mobilization mechanisms for synthetic biology toolkit development.

Community context altered both the breadth and identity of elements with evidence of excision. While some GIs showed similar excision rates in both contexts, many either had increased excision rates in community or excision was only detected during community growth. There was, however, a small subset of GIs (46.L and 49.R) that had decreased excision rates in the community. Together, these results support the conclusion that community context reshapes GI mobilization potential in ways not predictable from isolate studies alone. Furthermore, our study revealed a taxon-structured mobilome; that is, all GIs predicted in *Sinorhizobium, Neorhizobium*, and *Rhodococcus* had evidence of excision, while it was rare in *Variovorax*.

Phage-linked mobilization appears to dominate in MSC-1. Prophages, defective prophages, and defense GIs were always excised in MSC-1 isolates studied, while other types of GIs (ICE, metabolism, and others) were more variable. Widespread excision of phage-like GIs was unexpected given the low taxonomic redundancy within the MSC-1 community that would be predicted as suitable hosts for these phages to infect. This underscores a broader ecological role for prophages beyond traditional lysis, such as cargo delivery (39), metabolic remodeling (40), or promoting bacterial survival through phage-encoded defenses (41, 42). Further, the varied mobility of different types of GIs raises questions about how the nature of GI-encoded functions impacts overall microbiome function and by how much. While previous studies have demonstrated that HGT influences nutrient cycling, housekeeping functions, and signaling pathways, this likely just scratches the surface of the impact HGT can have on microbial community function and persistence (8, 43).

Understanding GI excision and integrase production can aid in the development of synthetic biology tools (44). GIs transfer large amounts of genetic cargo that can influence cell metabolism and phenotypes and, in many cases, stably persist in the genome for many generations. Understanding the mechanisms that GIs employ for transfer and stability could enable the development of sophisticated bioengineering tools. These platforms would allow for high efficiency introduction of genetic material to a target organism, even within complex communities, as well as long-term gene maintenance, expression and controllable excision. However, we must also consider biocontainment of both engineered organisms and genetic sequences of GIs, which can be spread within a community. We show that local conditions can activate distinct GI subsets from a large latent mobilome. Community composition and habitat conditions could compromise or enhance GI mobility which will impact features installed in these microbes (45). This context-dependent GI activation suggests that microbiome engineering strategies must be validated in relevant environmental contexts to ensure that the spread of engineered traits is limited (45). Rapid assessments that quantify transfer frequencies and host ranges under defined conditions and field contexts are needed to understand the basic biology and assess risks associated with deploying engineered microbes.

While GI excision rates quantified here provide a unique look at GI mobility in a model microbial community and the framework to better understand the impact of MGEs in communities, several considerations remain. 1) This work highlights the potential for GIs to mobilize under environmentally relevant conditions. Future work should aim to predict and track how these GIs may impact engineered traits. 2) GIs were abundantly excised under variable hydration levels, which were intended to mimic drought conditions (17), but hydration is only one of a myriad of environmental factor influencing soil microbiomes. Spatial structuring in microbial habitats and factors related to biofilm formation likely impact GI transfer within microbial communities (46, 47). Additional studies are necessary to understand how the spatial structure of microbiomes may impact the mobility and transfer partners of MGEs (48, 49). This is relevant not only to highly structured soil systems, but aquatic systems and host-associated microbiomes as well (17, 50, 51). 3) While we have shown these GIs excise from their specific host genome, it is unknown if they transfer to and integrate within the genomes of other members within the community. Future work could utilize sophisticated genetic toolkits for tracking the transfer of GIs between bacterial hosts (52, 53). Additionally, we uncovered many mobile prophages and defective prophages in our dataset that could potentially infect other members of MSC-1. Establishing community host ranges of these phages and understanding whether/when these phages pursue traditional lytic infection cycles, integrate as prophages, or reside as virocells could uncover control points for effectively co-opting phages for delivery of genetic cargo to aid in microbiome engineering. Unlocking these community-dependent control points could bridge the gap between lab models and native ecosystems, providing a foundational framework required to safely predict, manage and co-opt the genetic flux shaping soil microbiomes.

## STAR METHODS

Complete methods are listed in the supplemental material.

### Bioinformatic Genomic Island prediction and verification

TIGER and Islander (27, 28) were applied to MSC-1 isolate complete genomes and MSC-1 MAGs to precisely identify GIs using standard parameters. Tater was used within the TIGER pipeline to annotate each genomic island. DiMER was applied (https://github.com/sandialabs/DiMER) to further refine the functional predictions.

We applied TIGER using the flag -search IS to discover transposons within the MSC-1 isolate genomes. We searched the ISFinder database (54) using BLASTn to determine if any transposons predicted mapped to known IS elements.

### MMC induction of monocultures

MSC-1 isolates were separately cultured on solid R2A medium at the appropriate temperature (**Supplemental Table 5**). For induction experiments, liquid monocultures of each MSC-1 bacterial strain were grown in R2A broth to an OD_600_ of 0.5. Cultures were then induced with the addition of either 1 or 3 mg/mL of mitomycin C (**Supplemental Table 5)** and promptly returned to culture conditions. 1 ml of each induced culture was removed at 0, 1, 2, 3, 4, and 24 hours post mitomycin C induction. Sample supernatants were discarded and pellets were stored at −20°C prior to DNA extraction.

### Growth conditions for MSC-1 cocultures

Cocultures of MSC-1 with variable moisture saturation levels (100%, 50%, and 25%) were grown and gDNA, RNA, and protein samples were prepared previously (17).

### Identification of recombination events and GI excision

Recombination events across the genome were computationally identified with Juxtaposer (55), binned into 500 bp bins spanning the genome and visualized as histograms. Probes were generated for each *attP, attB, attL*, and *attR* site and reads were counted using attCt (55). To normalize the *att* site counts for **Figure 2**, we divided the *attB* and *attP* counts for each GI by the number of host genomes ((*attL*+*attB*)x(*attR*+*attB*))/2.

### Metatranscriptomics and Metaproteomic data analysis

Transcript activity was assessed by mapping metatranscriptomic reads using bbmap (https://archive.jgi.doe.gov/data-and-tools/software-tools/bbtools/) from MSC-1 incubation in a porous glass bead system with variable levels of moisture (17). Metatranscriptomic sequence alignment map (SAM) files were filtered to report hits of ≥98% identity and mapped to GI features using featureCounts (56). The resulting counts were then normalized in R to the total length of each respective gene to a Gene length corrected TMM value (GeTMM) (57). GeTMM values were then transformed to Z-scores.

Raw mass spectrometry data was searched against a targeted protein database of 231,411 translated proteins generated from the sequenced genomes of the MSC-1 consortium (39 members). After filtering out decoys and contaminants, peptide-spectrum matching resulted in 6,238 protein groups detected with non-zero intensity values.

## Acknowledgements

We thank Ryan McClure for his thoughtful discussions about this work and sharing the MSC-1 isolates with us. We thank Tagide deCarvalho at the University of Maryland Baltimore County Keith R. Porter Imaging Facility for TEM imaging MMC filtrates. We thank Elise Wilbourn for internal review of this manuscript.

Sandia National Laboratories is a multi-mission laboratory managed and operated by National Technology & Engineering Solutions of Sandia, LLC (NTESS), a wholly owned subsidiary of Honeywell International Inc., for the U.S. Department of Energy’s National Nuclear Security Administration (DOE/NNSA) under contract DE-NA0003525. This written work is authored by an employee of NTESS. The employee, not NTESS, owns the right, title and interest in and to the written work and is responsible for its contents. Any subjective views or opinions that might be expressed in the written work do not necessarily represent the views of the U.S. Government. The publisher acknowledges that the U.S. Government retains a non-exclusive, paid-up, irrevocable, world-wide license to publish or reproduce the published form of this written work or allow others to do so, for U.S. Government purposes. The DOE will provide public access to results of federally sponsored research in accordance with the DOE Public Access Plan.

## References

1. M. Juhas et al., Genomic islands: tools of bacterial horizontal gene transfer and evolution. FEMS Microbiol Rev 33, 376–393 (2009).

2. J. Hacker, E. Carniel, Ecological fitness, genomic islands and bacterial pathogenicity. A Darwinian view of the evolution of microbes. EMBO Rep 2, 376–381 (2001).

3. J. E. Silpe, O. P. Duddy, B. L. Bassler, Induction mechanisms and strategies underlying interprophage competition during polylysogeny. PLoS Pathog 19, e1011363 (2023).

4. D. J. Rankin, E. P. Rocha, S. P. Brown, What traits are carried on mobile genetic elements, and why? Heredity (Edinb) 106, 1–10 (2011).

5. C. M. Mageeney, G. Trubl, K. P. Williams, Improved Mobilome Delineation in Fragmented Genomes. Front Bioinform 2, 866850 (2022).

6. A. G. Kent, A. C. Vill, Q. Shi, M. J. Satlin, I. L. Brito, Widespread transfer of mobile antibiotic resistance genes within individual gut microbiomes revealed through bacterial Hi-C. Nat Commun 11, 4379 (2020).

7. X. Jiang, A. B. Hall, R. J. Xavier, E. J. Alm, Comprehensive analysis of chromosomal mobile genetic elements in the gut microbiome reveals phylum-level niche-adaptive gene pools. PLoS One 14, e0223680 (2019).

8. J. Guo et al., Mobile genetic elements shape microbial diversity and functions in thawing permafrost soils. Nat Microbiol 11, 1800–1814 (2026).

9. A. P. Camargo et al., Identification of mobile genetic elements with geNomad. Nat Biotechnol 42, 1303–1312 (2024).

10. L. S. Frost, R. Leplae, A. O. Summers, A. Toussaint, Mobile genetic elements: the agents of open source evolution. Nat Rev Microbiol 3, 722–732 (2005).

11. L. A. Shakes, H. M. Wolf, D. C. Norford, D. J. Grant, P. K. Chatterjee, Harnessing mobile genetic elements to explore gene regulation. Mob Genet Elements 4, e29759 (2014).

12. A. Chaaban, R. Sleem, J. Santina, M. Rima, J. N. Ibrahim, Exploring transposable elements: new horizons in cancer diagnostics and therapeutics. Mob DNA 16, 28 (2025).

13. Y. Li et al., Roles of mobile genetic elements and biosynthetic gene clusters in environmental adaptation of acidophilic archaeon Ferroplasma to extreme polluted environments. Front Microbiol 16, 1654373 (2025).

14. Z. Saati-Santamaría, A. Flores, I. Canosa, P. García-Fraile, Horizontal Gene Transfer and Genome Rearrangements Shape Bacterial Adaptation for Bioremediation. Environ Microbiol 28, e70374 (2026).

15. R. McClure et al., Interaction Networks Are Driven by Community-Responsive Phenotypes in a Chitin-Degrading Consortium of Soil Microbes. mSystems 7, e0037222 (2022).

16. R. McClure et al., Development and Analysis of a Stable, Reduced Complexity Model Soil Microbiome. Front Microbiol 11, 1987 (2020).

17. J. Rodríguez-Ramos et al., Environmental matrix and moisture influence soil microbial phenotypes in a simplified porous media incubation. mSystems 10, e0161624 (2025).

18. A. E. Shikov, Y. V. Malovichko, A. A. Nizhnikov, K. S. Antonets, Current Methods for Recombination Detection in Bacteria. Int J Mol Sci 23 (2022).

19. L. A. Matthews, L. A. Simmons, Bacterial nonhomologous end joining requires teamwork. J Bacteriol 196, 3363–3365 (2014).

20. D. Liao, Gene conversion drives within genic sequences: concerted evolution of ribosomal RNA genes in bacteria and archaea. J Mol Evol 51, 305–317 (2000).

21. R. P. Anderson, J. R. Roth, Tandem genetic duplications in phage and bacteria. Annu Rev Microbiol 31, 473–505 (1977).

22. C. W. Hill, Large genomic sequence repetitions in bacteria: lessons from rRNA operons and Rhs elements. Res Microbiol 150, 665–674 (1999).

23. C. W. Hill, B. W. Harnish, Inversions between ribosomal RNA genes of Escherichia coli. Proc Natl Acad Sci U S A 78, 7069–7072 (1981).

24. P. Anderson, J. Roth, Spontaneous tandem genetic duplications in Salmonella typhimurium arise by unequal recombination between rRNA (rrn) cistrons. Proc Natl Acad Sci U S A 78, 3113–3117 (1981).

25. P. Siguier, E. Gourbeyre, A. Varani, B. Ton-Hoang, M. Chandler, Everyman’s Guide to Bacterial Insertion Sequences. Microbiol Spectr 3, Mdna3-0030-2014 (2015).

26. E. Nzabarushimana, H. Tang, Insertion sequence elements-mediated structural variations in bacterial genomes. Mob DNA 9, 29 (2018).

27. C. M. Hudson, B. Y. Lau, K. P. Williams, Islander: a database of precisely mapped genomic islands in tRNA and tmRNA genes. Nucleic Acids Res 43, D48–53 (2015).

28. C. M. Mageeney et al., New candidates for regulated gene integrity revealed through precise mapping of integrative genetic elements. Nucleic Acids Res 48, 4052–4065 (2020).

29. C. Mageeney et al., Mycobacteriophage Marvin: a new singleton phage with an unusual genome organization. J Virol 86, 4762–4775 (2012).

30. A. Fossati et al., Next-generation proteomics for quantitative Jumbophage-bacteria interaction mapping. Nat Commun 14, 5156 (2023).

31. R. A. Frampton et al., Genome, Proteome and Structure of a T7-Like Bacteriophage of the Kiwifruit Canker Phytopathogen Pseudomonas syringae pv. actinidiae. Viruses 7, 3361–3379 (2015).

32. J. W. Wu, X. L. Chen, Extracellular metalloproteases from bacteria. Appl Microbiol Biotechnol 92, 253–262 (2011).

33. T. Hackl et al., Novel integrative elements and genomic plasticity in ocean ecosystems. Cell 186, 47–62.e16 (2023).

34. A. Ruf et al., Extracellular Vesicles From Xylella fastidiosa Carry sRNAs and Genomic Islands, Suggesting Roles in Recipient Cells. J Extracell Vesicles 14, e70102 (2025).

35. S. Takano et al., Enrichment of horizontally transferred gene clusters in bacterial extracellular vesicles via non lytic mechanisms. Isme j 19 (2025).

36. S. J. Biller et al., Distinct horizontal gene transfer potential of extracellular vesicles versus viral-like particles in marine habitats. Nat Commun 16, 2126 (2025).

37. S. L. Yu, C. M. Mageeney, F. Shormin, N. Ghaffari, K. P. Williams, Speeding genomic island discovery through systematic design of reference database composition. PLoS One 19, e0298641 (2024).

38. T. Nogueira et al., Horizontal gene transfer of the secretome drives the evolution of bacterial cooperation and virulence. Curr Biol 19, 1683–1691 (2009).

39. P. Forterre, The virocell concept and environmental microbiology. Isme j 7, 233–236 (2013).

40. C. Howard-Varona et al., Phage-specific metabolic reprogramming of virocells. Isme j 14, 881–895 (2020).

41. J. Bondy-Denomy et al., Prophages mediate defense against phage infection through diverse mechanisms. Isme j 10, 2854–2866 (2016).

42. R. M. Dedrick et al., Prophage-mediated defence against viral attack and viral counter-defence. Nat Microbiol 2, 16251 (2017).

43. A. T. Bisesi, R. P. Carlson, L. Cotner, W. R. Harcombe, Metabolic remodeling of microorganisms by mobile genetic elements alters mutualistic community composition. mSystems 10, e0014425 (2025).

44. H. M. McClain et al., Integrase-On-Demand: bioprospecting integrases for targeted genomic insertion of genetic cargo. Nucleic Acids Res 54 (2026).

45. J. E. McDermott et al., Describing the persistence landscape for introducing microbes into complex communities. Appl Environ Microbiol 10.1128/aem.00930-26, e0093026 (2026).

46. C. C. Goller, T. Romeo, Environmental influences on biofilm development. Curr Top Microbiol Immunol 322, 37–66 (2008).

47. Y. Joshi et al., Unearthing and harnessing the role of microbial biofilms in soil ecosystems: Structure, function and potential for enhancing soil integrity and climate mitigation. Agriculture Communications, 100133 (2026).

48. V. Warrier et al., Interplay of Spatial Structure and Interactions in Microbial Communities. Environ Microbiol 28, e70262 (2026).

49. M. Bäcker et al., Spatial structure: shaping the ecology and evolution of microbial communities. FEMS Microbiol Rev 50 (2026).

50. B. Grodner et al., Spatial mapping of mobile genetic elements and their bacterial hosts in complex microbiomes. Nat Microbiol 9, 2262–2277 (2024).

51. I. Ntekas et al., Spatial transcriptomics maps host-gut microbiome biogeography at high resolution. Nat Microbiol 11, 1193–1204 (2026).

52. V. Partanen et al., Use of sequence barcodes for tracking horizontal gene transfer of antimicrobial resistance genes in a microbial community. ISME Commun 5, ycaf113 (2025).

53. P. B. Kalvapalle et al., Information storage across a microbial community using universal RNA barcoding. Nat Biotechnol 44, 269–276 (2026).

54. P. Siguier, J. Perochon, L. Lestrade, J. Mahillon, M. Chandler, ISfinder: the reference centre for bacterial insertion sequences. Nucleic Acids Res 34, D32–36 (2006).

55. J. S. Schoeniger, C. M. Hudson, Z. W. Bent, A. Sinha, K. P. Williams, Experimental single-strain mobilomics reveals events that shape pathogen emergence. Nucleic Acids Res 44, 6830–6839 (2016).

56. Y. Liao, G. K. Smyth, W. Shi, featureCounts: an efficient general purpose program for assigning sequence reads to genomic features. Bioinformatics 30, 923–930 (2014).

57. M. Smid et al., Gene length corrected trimmed mean of M-values (GeTMM) processing of RNA-seq data performs similarly in intersample analyses while improving intrasample comparisons. BMC Bioinformatics 19, 236 (2018).

58. J. P. Quast, D. Schuster, P. Picotti, protti: an R package for comprehensive data analysis of peptide- and protein-centric bottom-up proteomics data. Bioinform Adv 2, vbab041 (2022).

59. S. Koren et al., Canu: scalable and accurate long-read assembly via adaptive k-mer weighting and repeat separation. Genome Res 27, 722–736 (2017).

60. C. Jain, R. L. Rodriguez, A. M. Phillippy, K. T. Konstantinidis, S. Aluru, High throughput ANI analysis of 90K prokaryotic genomes reveals clear species boundaries. Nat Commun 9, 5114 (2018).

61. W. McKinney, Data structures for statistical computing in Python. scipy 445, 51–56 (2010).

62. M. L. Waskom, Seaborn: statistical data visualization. Journal of open source software 6, 3021 (2021).

63. J. D. Hunter, Matplotlib: A 2D graphics environment. Computing in science & engineering 9, 90–95 (2007).

64. P. Shannon et al., Cytoscape: a software environment for integrated models of biomolecular interaction networks. Genome Res 13, 2498–2504 (2003).

65. C. L. M. Gilchrist, Y. H. Chooi, clinker & clustermap.js: automatic generation of gene cluster comparison figures. Bioinformatics 37, 2473–2475 (2021).

66. J. Rodríguez-Ramos et al., Environmental matrix and moisture are key determinants of microbial phenotypes expressed in a reduced complexity soil-analog. [Data Set] PNNL DataHub.

67. T. Seemann, Prokka: rapid prokaryotic genome annotation. Bioinformatics 30, 2068–2069 (2014).

68. J. Cox, M. Mann, MaxQuant enables high peptide identification rates, individualized p.p.b.-range mass accuracies and proteome-wide protein quantification. Nat Biotechnol 26, 1367–1372 (2008).

69. S. Tyanova, T. Temu, J. Cox, The MaxQuant computational platform for mass spectrometry-based shotgun proteomics. Nat Protoc 11, 2301–2319 (2016).

70. K. S. Hofmockel, Environmental matrix and moisture are key determinants of microbial phenotypes expressed in a reduced complexity soil-analog. [Data set] MassIVE MSV000103079.

