## Supplemental Figure for "Mobile genetic elements are active and responsive to community context in model microbial consortium"

### Supplemental Information

#### Expanded Materials and Methods

##### Reference genome generation

We constructed new reference genomes for *Streptomyces sp*. MSC-1_001, *Variovorax beijingensis* MSC-1_012, *Sinorhizobium meliloti* MSC-1_014, and *Rhodococcus sp.* MSC-1_016 to ensure complete genomes with a single contig were available for all MSC-1 isolate members. Liquid monocultures of each MSC-1 isolate strain were grown independently in R2A broth to an $OD_{600}$ of 1.0. Cell pellets from 1 ml of each culture were stored at -20°C prior to DNA extraction. DNA was extracted using the PacBio CBB Nanobind kit (PacBio catalog # 102-301-900) kit. Genomic DNA was shared and prepared into SMRTbell libraries using the SMRTbell Express Template Prep Kit 3.2 (PacBio, catalog # 602826) following the manufacturer's protocol. Sequencing was performed on the PacBio Sequel IIe System according to the manufacturer's recommendations.

Canu (v2.2) (59) was used to assemble PacBio long read sequencing data for *Variovorax beijingensis* MSC-1_012 and *Sinorhizobium meliloti* MSC-1_014. Standard parameters were used except for the genomeSize parameter, which was determined by the genome size of the short-read assembly (i.e genomeSize for Variovorax was set to 6.7m).

Whole genome sequencing of *Streptomyces sp*. MSC-1_001 and *Rhodococcus sp.* MSC-1_016 was performed by Plasmidsaurus using Oxford Nanopore technology with custom analysis and annotation.

##### Bioinformatic genomic island prediction and verification

The *Sphingopyxis* GI 212.S was manually curated because TIGER has an upper size limit of 200kbp. We noticed during our analysis that the GI called 193.Y-Int (1609616-1802501) has no recombination activity. We did see high levels of activity in transcriptomics and many proteins in the proteomics dataset from this GI. To determine the precise end points, we mapped the integrase sequence to our large database of integrases (44) to find the closest match. We then searched our genome for the *attL* and *attR* sequences corresponding to the reference integrase. Once the end point was identified, we blasted each end to ensure the identity block within the *att* site was fully captured. These refined coordinates are listed in **Supplemental Table 2.**

We used fastANI (60) to compare all MSC-1 isolates and MSC-1 community member genomes. Python packages pandas v3.0.0 (61), seaborn v0.13.2 (62), and matplotlib v3.10.8 (63) were used to convert the output and generate the ANI heatmap (**Supplemental Figure 13**).

We clustered the MSC-1 GIs using amino acid identity using the AAI_cluster pipeline with the following parameters for genus level clustering (--min_percent_shared 20 --min_num_shared 16 --min_aai 40) (https://github.com/snayfach/MGV/blob/master/aai_cluster/). The clusters were visualized with cytoscape (64).

Nucleotide comparative genomic maps were created using custom python scripts calling pandas v3.0.0 (61) and matplotlib v3.10.8 (63) for nucleotide alignment which required BLASTn outputs between the two GIs compared. Amino acid comparative genomics maps were created using clinker (65) with gff inputs of each GI.

##### Clustering and read attribution of GIs

To confidently attribute excision events within consortium members, we used average nucleotide identity (ANI) to determine if the MSC-1 isolates shared nucleotide similarity with other members of the

MSC-1 community that could lead to ambiguous mapping of sequences (**Supplemental Figure 13**). As expected, each isolate had 100% ANI over the entire genome with itself (**Supplemental Figure 13A**).

The remaining seven isolates share genome nucleotide similarity with other MSC-1 community members (**Supplemental Figure 13B-H**). *Variovorax* shares nucleotide similarity with six MSC-1 members but with a single exception has low alignment fraction (AF) values (<40%), enabling confident assignment of reads to this genome (**Supplemental Figure 13D**). *Sphingopyxis*, *Dyadobacter*, and *Streptomyces* share minimal nucleotide identity with MSC-1 members (ANI value <80% and alignment <50%) (**Supplemental Figure 13C, E, and F**). The final three MSC-1 isolates, *Sinorhizobium*, *Neorhizobium*, and *Ensifer* share considerable similarity with each other and three additional members of MSC-1 (**Supplemental Figure 13B, G-H**), impacting our ability to confidently attribute reads to only a single member.

We sought to accurately assign transcript and protein-level expression to only the expected GI. Comparative genomics maps were created with blast and clinker to visualize the locations of the shared content (**Supplemental Figure 3**).

**Whole genome deep-sequencing**

DNA was isolated from MMC-treated cell pellets using the Qiagen DNeasy Blood and Tissue Kit (Qiagen, catalog # 69506). DNA libraries were prepared with the Illumina DNA Prep, (M) Tagmentation library prep kit using Illumina DNA/RNA UD Indexes Set B; following the manufacturer recommended protocol. The samples were multiplexed and sequenced on a NextSeq 2000 and run for 160 cycles using NextSeq 2000 P3 XLEAP-SBS reagent kit (Illumina, catalog # 20100989). To repeat sequencing runs for samples with low depth or varying MMC concentrations, purified gDNA was sequenced by Azenta.

**Identification of recombination events and GI excision**

To quantify GI excision events, we generated DNA probes sequences that matched the sequences of the *attL* and *attR*, and the predicted *attB* and *attP* from TIGER outputs (**Supplemental Table 2**). The GI boundaries of 212.S in *Sphingopyxis* was incorrectly identified and thus we manually identified the full island and generated corresponding *att* site probes.

Using Juxtaposer (55), the number of sequences containing each of the *att* probe sequences within the whole genome deep sequencing or MSC-1 metagenomic raw reads (66) were counted. To normalize by the number of cells within the population, *attB* and *attP* counts were divided by the number of cells in the population (n = ((*attL*+*attB*) + (*attR*+*attB*))/2).

**Metatranscriptomics data analysis**

Transcript activity was assessed by mapping metatranscriptomic reads from an incubation of microbial consortia in a glass bead system (66). Briefly, we use a reduced-complexity microbial consortium grown in a glass bead porous medium amended with chitin to test the effects of moisture and a structural matrix on microbial phenotypes (17). In total, 25 metatranscriptomic read samples to a reference database of the Model Soil Consortia (MSC-1) using bbmap from the bbtools suite (https://archive.jgi.doe.gov/data-and-tools/software-tools/bbtools/. To remove ribosomal RNA (rRNA) contamination from metatranscriptomes, reads were mapped to a database of rRNA from our assembled metagenomes identified using barrnap (https://github.com/tseemann/barrnap) with flags ambiguous = all and perfectmode = t. Reads that perfectly mapped to rRNA were then removed, and subsequent cleaned reads were used. After initial mapping, sequence alignment map (SAM) files were then filtered to report hits of ≥98% identity (minidfilter = 0.98) using reformat.sh from bbsuite (https://sourceforge.net/projects/bbmap/files/). Gff files for the genes from the MSC-1 database were generated with our tater annotation pipeline, which calls Prokka v. 1.11 (67) for open reading frame prediction, uses tFind, and rFind software for RNA prediction, and prodigal for functional predictions (https://github.com/sandialabs/TIGER) (28). Metatranscriptomic mapping SAM files were processed with featureCounts (56) using the subread package from Bioconda (https://bioconda.github.io/recipes/subread/README.html) along with the flags -t rrna, trna, CDS, -s 2 (reversely stranded), -M (multi-map), and -p (paired). The resulting counts were then normalized in R to the total length of each respective gene and subsequently converted to transcripts per million (TPM) according to the code in Additional File 4 of the geTMM manuscript (57).

Raw TPM were transformed to Z-score to normal across all GIs or all genes in each GI separately. The TPM was log2 normalized. The average and standard deviation were calculated for the entire dataset being compared (full GI or all genes). The Z-score was calculated using the following equation:

Z-score = L2 Normalized TPM – Average Log2 TPM/ Standard deviation of Log2 TPM

**Metaproteomics data analysis**

Raw MS data were searched against a targeted protein database of 231,411 translated proteins generated from the sequenced genomes of the MSC-1 consortium (39 members) with MSC-2 consortium (8 members) nested therein. The database included 132 common protein contaminants from the cRAP database (https://www.thegpm.org/crap/), as well as multiple trypsin and human keratin sequences. MaxQuant (v2.5.1.0; (68, 69)) was run in label-free quantification (LFQ) and Match Between Runs (MBR) modes for peptide-spectrum matching, with 20ppm parent ion mass tolerance, variable methionine oxidation, and considering partially tryptic peptides. Cysteine carbamidomethylation was included as a static modification. Matched peptides were controlled to a 1% false discovery rate at the peptide level using the target-decoy method. 245,292 peptide-spectrum matches were made from searching 799,151 total spectra, representing 46,133 unique peptides associated with 10,298 unique proteins. Raw and processed proteomics data have been deposited to the ProteomeXchange Consortium via the MassIVE repository (https://massive.ucsd.edu/ProteoSAFe) under accession number MSV000103079 (70).

After filtering out decoys and contaminants, peptide-spectrum matching resulted in 6,238 protein groups detected (proteins that cannot be unambiguously identified by unique peptides) with non-zero intensity values. We excluded protein groups represented by 6 or more proteins from further analysis since these would be difficult to confidently assign to a specific member of the consortium (n = 5,684 protein groups analyzed). Two samples were removed from analysis due to poor quality, both from the lowest moisture bead treatment (msc1_5_global_rep1_14 -- only 68 proteins; msc1_5_global_rep3_16 -- only 15 proteins). Between 2277 and 3737 protein groups were detected per sample. Protein group LFQ intensity values were log2-transformed and median normalized using the R package protti (58). Resulting data were filtered to only proteins identified on annotated genomic islands for further analysis (n = 144 protein groups). For heatmaps, intensity values were z-score scaled for visualization, excluding missing values.

**DNA pileups**

Pileups from monoculture DNA sequencing were created by aligning cutadapt trimmed reads to the reference genomes using bowtie2. Non-uniquely mapped reads were then removed with sed. Data was then sorted and indexed with samtools, and strand coverage was determined with bedtools genomecov.

**Transmission electron microscopy imaging of phage particles**

Monocultures of each MSC-1 member were grown in 4 ml of R2A medium at 30 °C. When an $OD_{600}$ of 0.5 was reached, cultures were treated with MMC at appropriate concentration (Table 1) and returned to growth conditions. After 4 hours, cultures were centrifuged, and supernatants were filtered through a 0.2 mM filter. Filtered supernatants were stored at 4 °C. Supernatants were concentrated by centrifuging at 4000 x G for 4 h at 4 °C and carefully decanting ¾ of the total volume from the surface of the liquid. Samples were imaged on a Hitachi HT7800 TEM at University of Maryland Baltimore County.

**Supplemental Tables**

All provided in supplemental excel spreadsheet.

**Supplemental Table 1. Recombination events per 1 million sequencing reads.** Recombination events per million are shown at each timepoint for the eight MSC-1 members following MMC induction (hours).

**Supplemental Table 2. Table of MSC-1 isolate genomic island predictions.** We list the strain of each isolate, genomic locations (contig, left and right coordinates, and direction), GI name (length in kb.insertion site), GI length, GI type, mobility observations in isolate and co-culture conditions, information about the integrase gene(s), support scores, and *attB*, *attP*, *attL*, and *attR* probes used for analysis.

**Supplemental Table 3. Table of MSC-1 genomic island predictions.** We list the remaining predicted genomic islands for all MSC-1 members using the same format listed for the isolates.

**Supplemental Table 4. MSC-1 isolate transposons.** We list the strain of each isolate, genomic location, transposon name (length in kb.insertion site), length, score, transposase gene, and what known IS element it was mapped to.

**Supplemental Table 5. MSC-1 isolate growth and induction conditions.** The 8 isolates from the MSC-1 consortium, their designation, and the temperature in which the bacteria were grown, and the MMC concentration added for MMC induction experiments.

#### Supplemental Figures

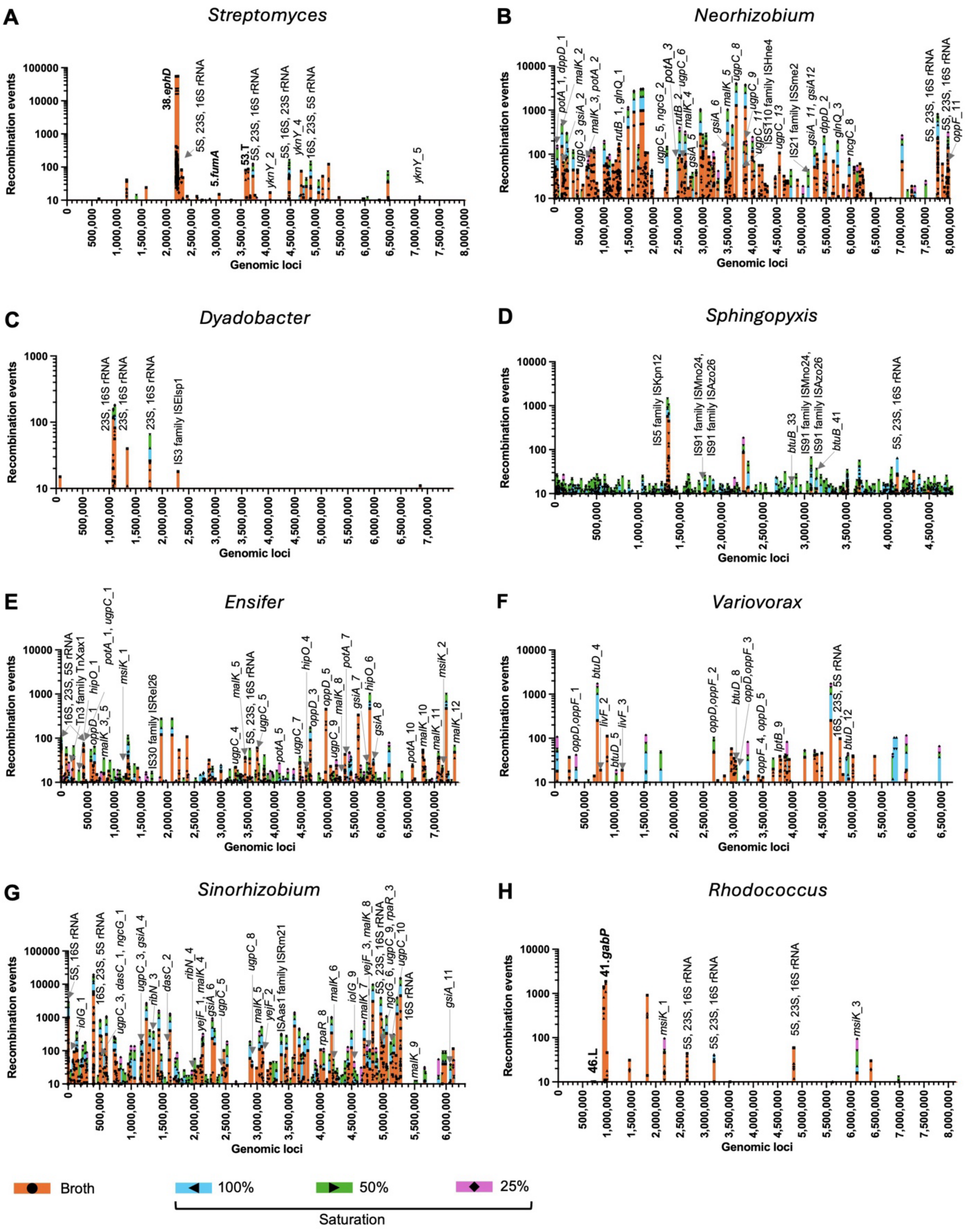

**Supplemental Figure 1. DNA recombination events with MSC-1 members grown as an MSC-1 community under different saturation conditions.** The number of DNA recombination events identified by high-depth, short-read sequencing were sorted into 500 bp bins spanning the genome. Histograms (A-H) were produced for (A) *Streptomyces*, (B) *Neorhizobium*, (C) *Dyadobacter*, (D) *Sphingopyxis*, (E) *Ensifer*, (F) *Variovorax*, (G) *Sinorhizobium*, and (H) *Rhodococcus*. IS-element and GI recombination events are labeled with the IS-elements name and family and GI name respectively. Suspected homologous recombination events are labeled with the homologous genes at the boundaries. All growth conditions (growth in liquid medium (Broth) or different saturation levels on porous glass beads) are plotted (key below).

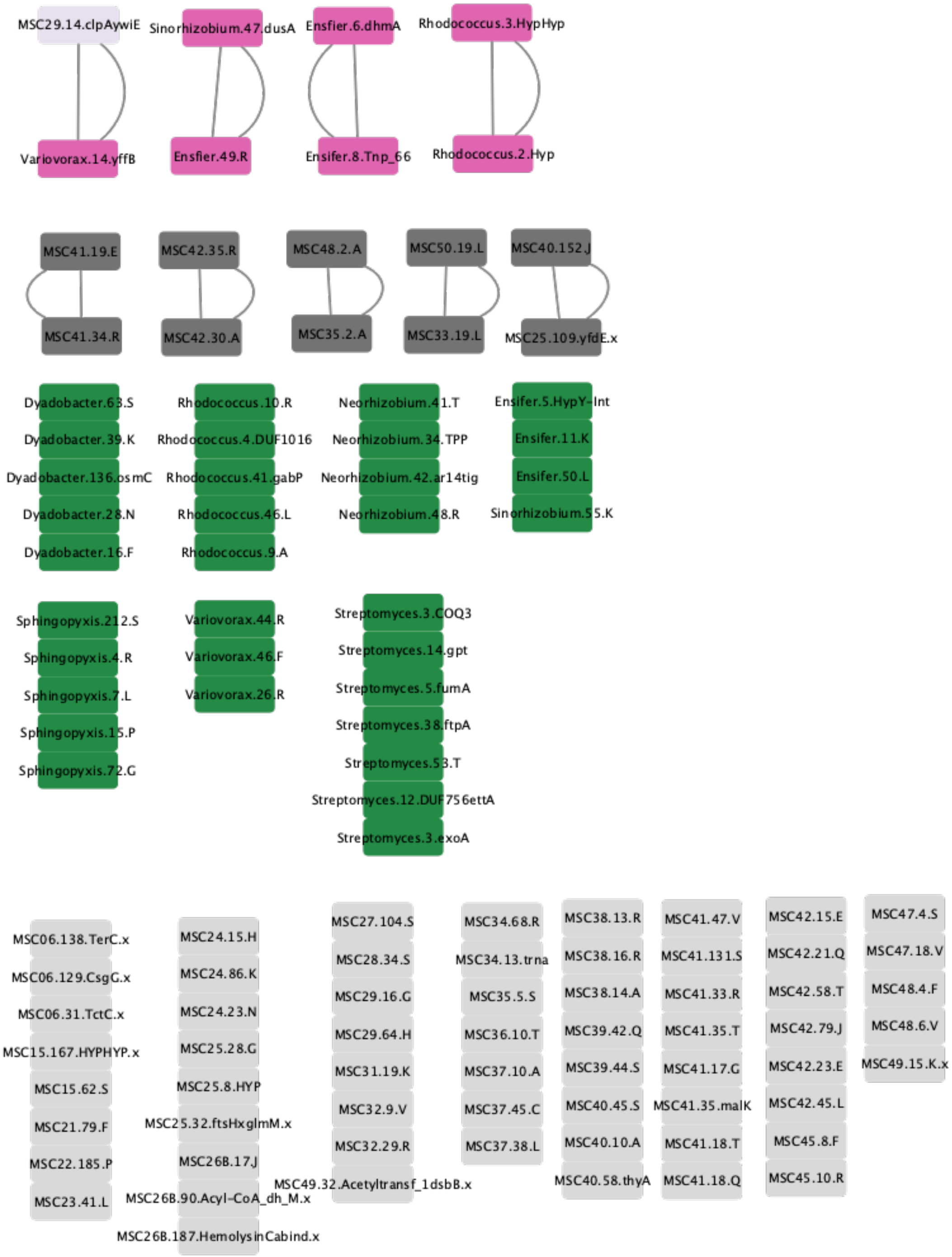


**Supplemental Figure 2. Cluster Network for MSC-1 GIs.** GIs were clustered using amino acid identity (see methods). Unique MSC-1 GIs not found in the MSC-1 isolates are colored light grey. MSC-1 isolate GIs that are unique are dark green. Clusters containing MSC-1 isolate GIs are colored in dark pink (MSC-1 isolates) and light pink (MSC-1 MAGs). Clusters containing no MSC-1 isolate GIs are colored in dark grey.

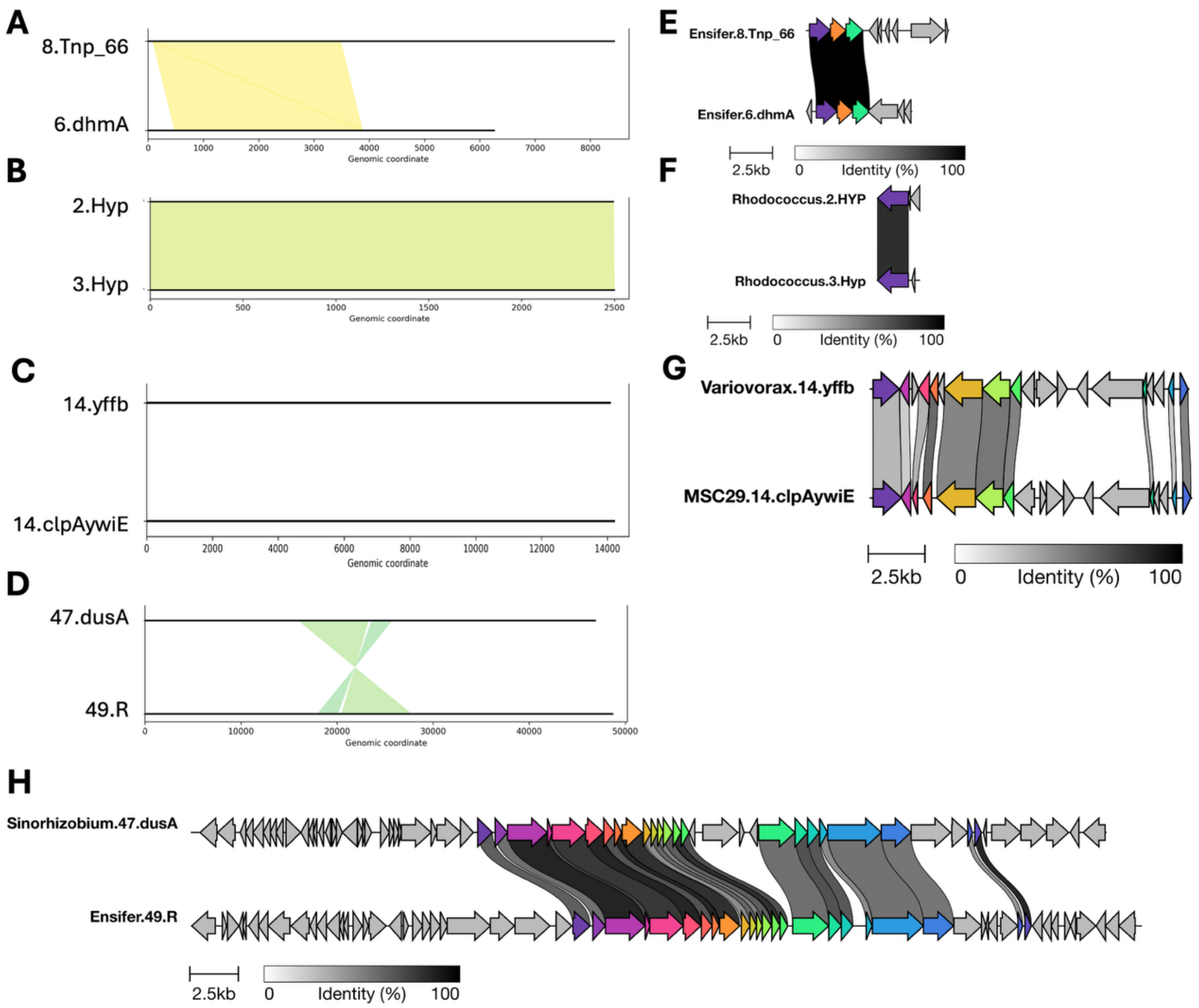


**Supplemental Figure 3. Comparative genomics maps of MSC-1 clusters.** (**A-D**) Nucleotide alignments are shown for (**A**) *Ensifer* GIs, (**B**) <u>*Rhodococcus*</u> GIs, (**C**) *Variovorax* 14.yffB and MSC-19 14.clpAywiE, (**D**) *Sinorhizobium* 47.dusA and *Ensifer* 49.R. (**E-H**) Amino acid alignments are shown with grey scale key representing percent identity and scale bars. Genes that are similar (**E**) *Ensifer* GIs, (**F**) *Rhodococcus* GIs, (**G**) *Variovorax* 14.yffB and MSC-19 14.clpAywiE, (**H**) *Sinorhizobium* 47.dusA and *Ensifer* 49.R.

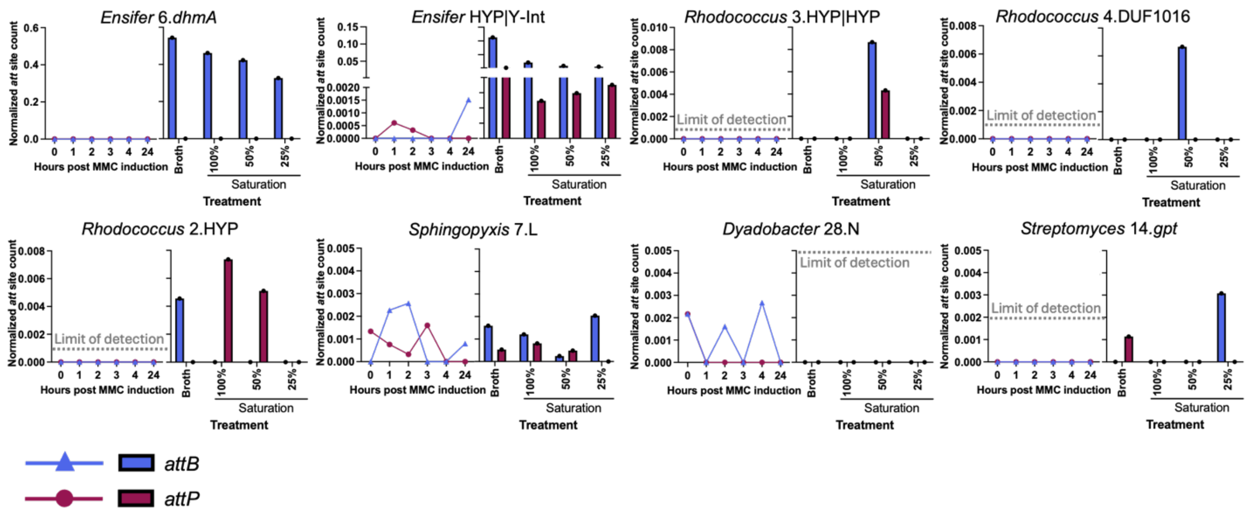


**Supplemental Figure 4. Normalized *attP* and *attB* of GIs unable to be typed.** Normalized *attP* (red circles/bars) and *attB* (blue triangles/bars) counts for GIs under monoculture MMC induction (left, line graph) and coculture growth with variable medium saturation (right, grouped bar graph) for all induced genomic islands classified as “other”. Key at bottom left. Cocultures grown in liquid medium are listed as “Broth".

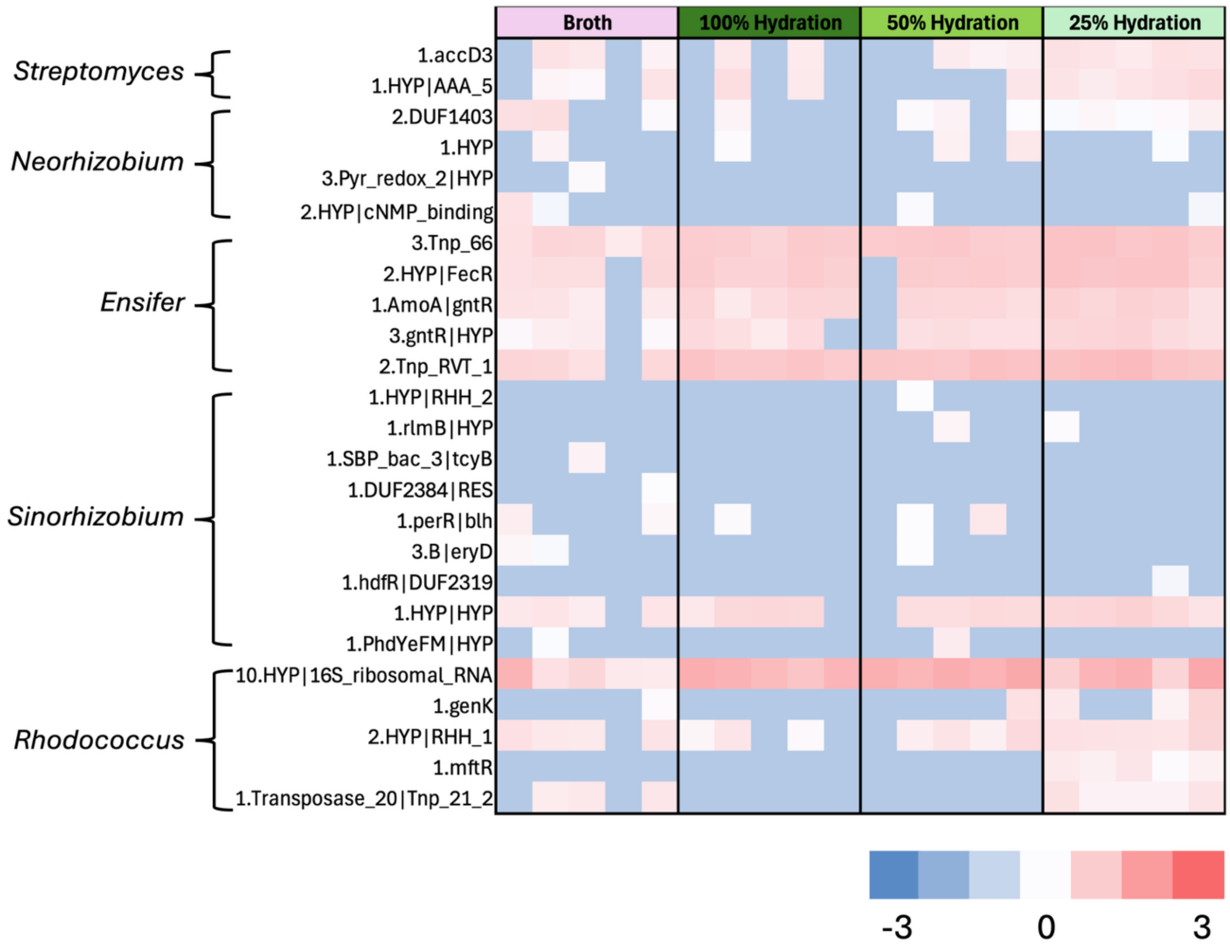


**Supplemental Figure 5. Z-score normalized TPM for predicted transposons.** The strain and transposon name are listed on the left side, and the color key is on the bottom right.

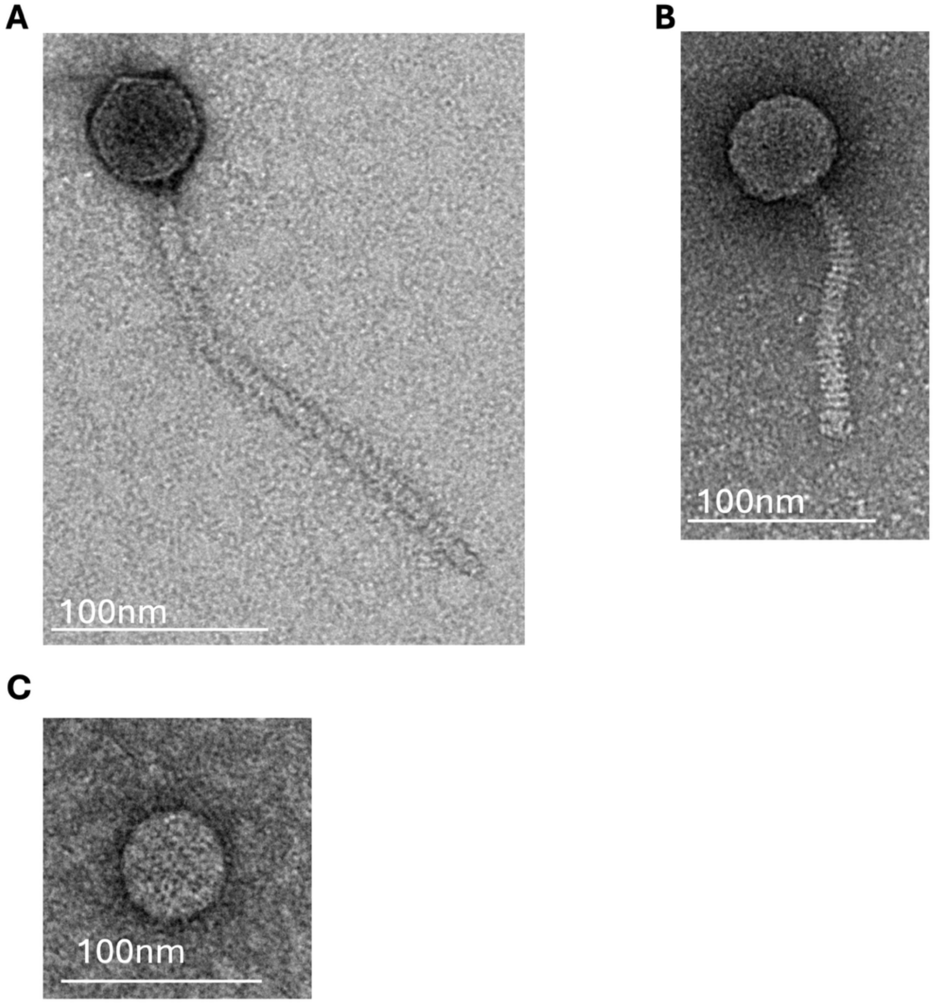


**Supplemental Figure 6. TEM images of MMC-induced isolate filtrates**. The scale bar is listed with each image. (**A**) Image of phage induced from *Rhodococcus* sp. MSC1_016. (**B**) Image of phage isolated from *Streptomyces* sp. MSC1_001. (**C**) Image of vesicle found after induction of *Sphingopyxis* sp. MSC1_008

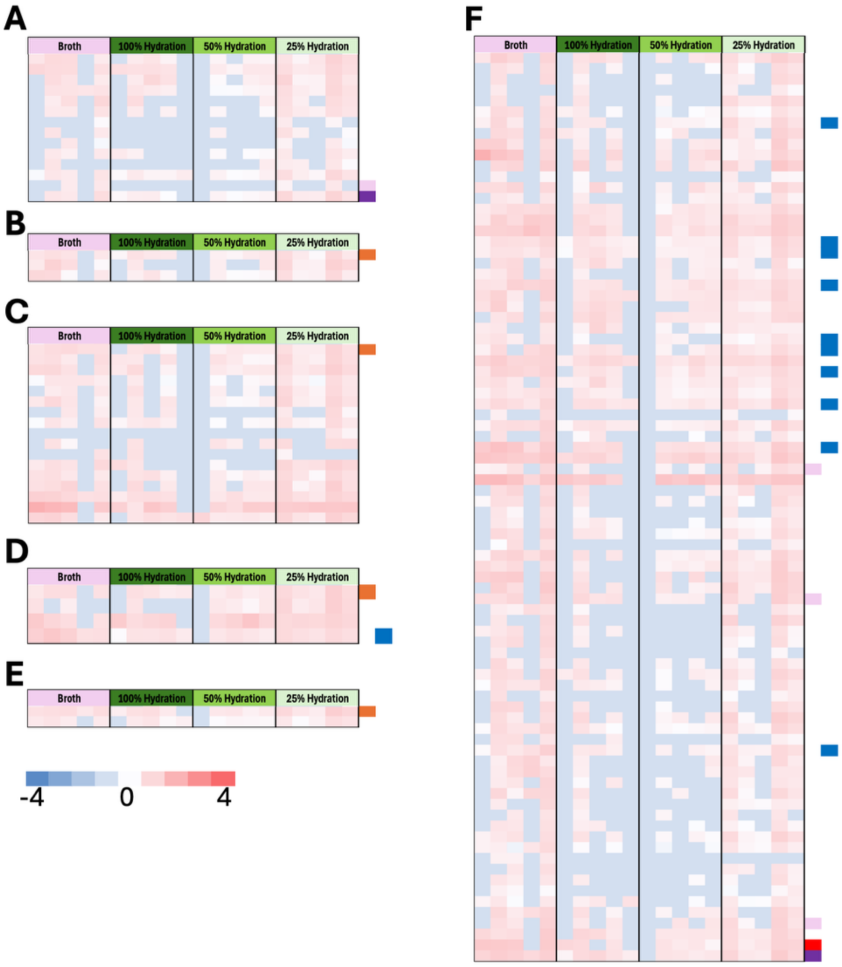


**Supplemental Figure 7. Z-score normalized TPM heatmaps of predicted genes in *Streptomyces* sp. MSC1_001 GIs**. The condition is listed across the top of each heatmap, and the color key is in the lower left corner. (**A**) 12.DUF756|ettA, (**B**) 3.exoA, (**C**) 14.gpt, (**D**) 5.fumA, (**E**)3.COQ3, (**F**) 53.T.

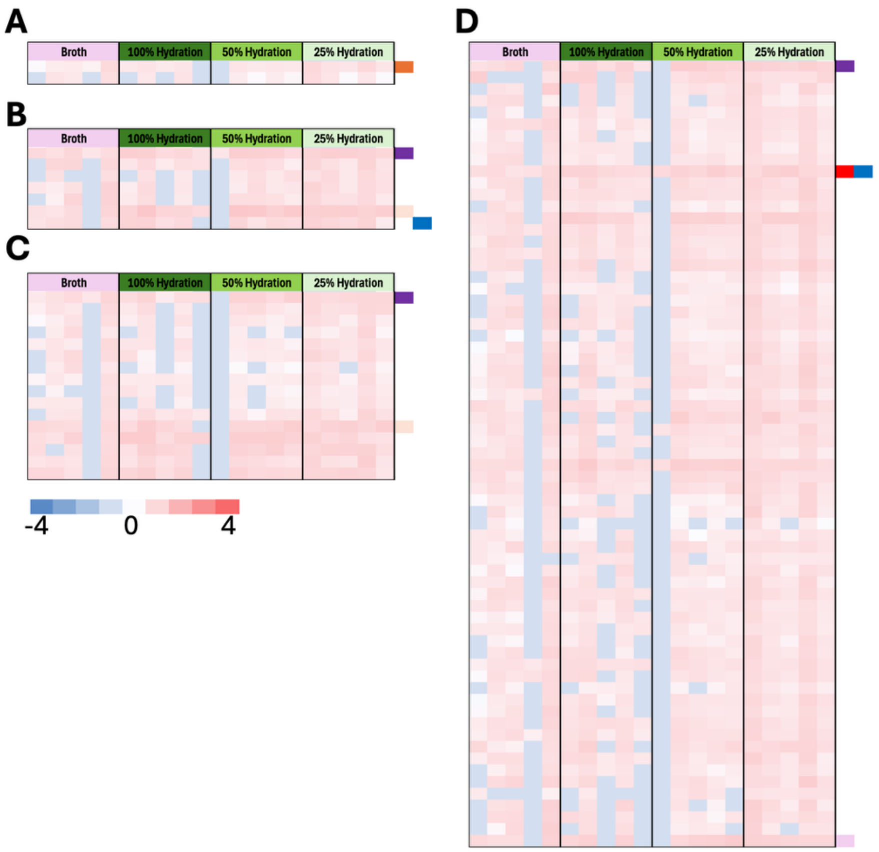


**Supplemental Figure 8. Heat maps of Z-score normalized TPM of genes in *Sphingopyxis* sp. MSC1_008 GIs**. The conditions are listed across the top and the color key is in the lower left corner. (**A**) 4.R, (**B**) 7.L, (**C**) 15.P, (**D**) 72.G.

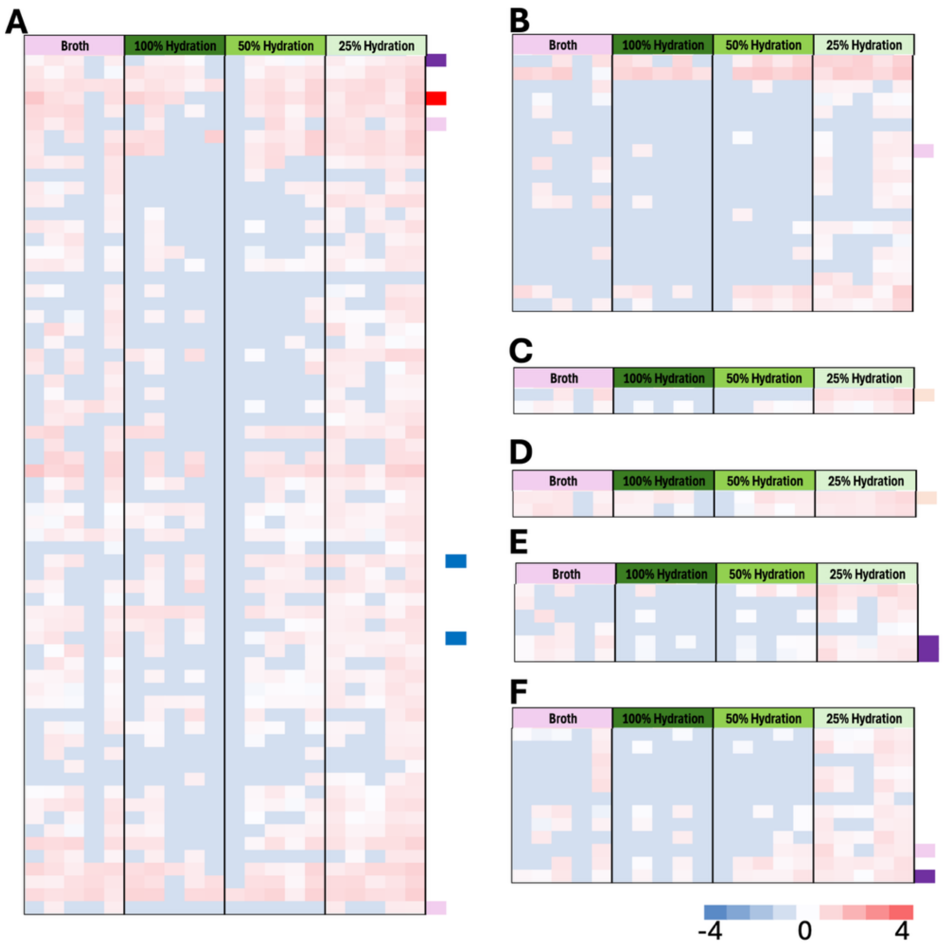


**Supplemental Figure 9. Z-score normalized TPM heatmap for genes in *Rhodococcus* sp. MSC1_016 GIs**. Conditions and color key are listed. (**A**) 46.L, (**B**) 10.R, (**C**) 3.Hyp|Hyp, (**D**) 2.Hyp, (**E**) 4.DUF1016, (**F**) 9.A.

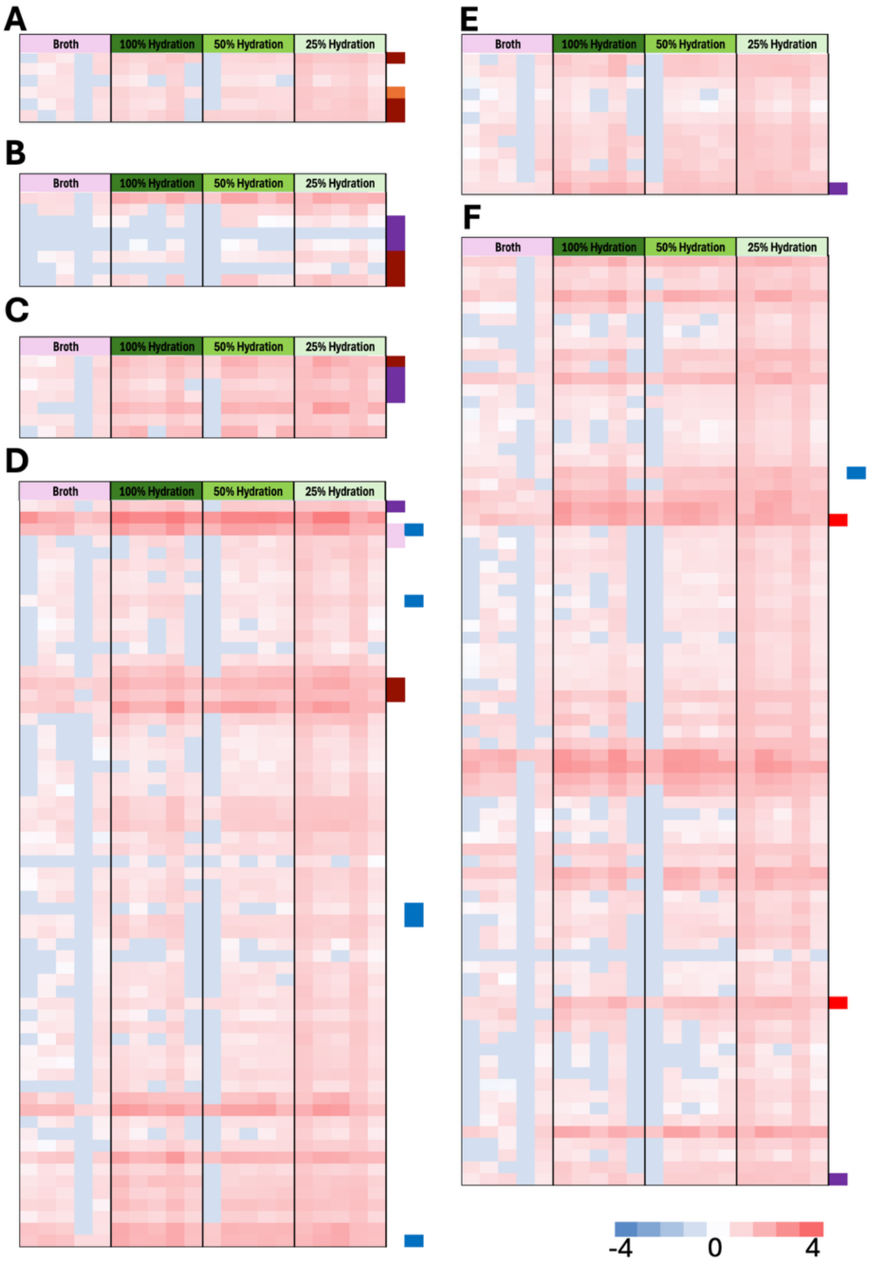


**Supplemental Figure 10. Z-score normalized TPM heatmaps for genes predicted in *Ensifer adhaerens* MSC1_011**. The conditions are listed across the top, and the color key is in the lower right corner. (**A**) 5.Hyp|Y-Int, (**B**) 6.dhmA, (**C**) 8.Tnp_66, (**D**) 49.R, (**E**) 11.K, (**F**) 50.L.

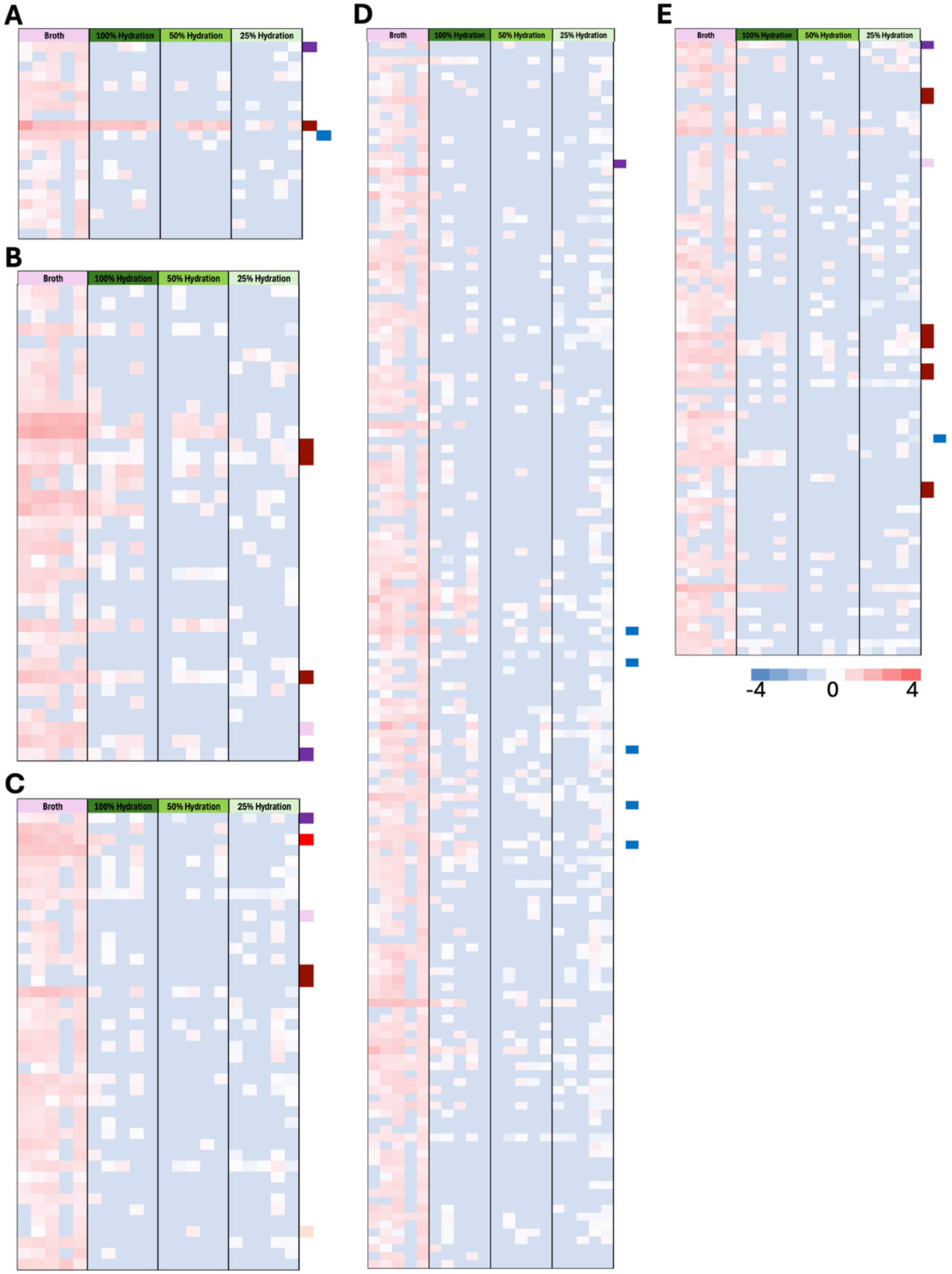


**Supplemental Figure 11. TPM normalized by Z-score for genes in *Dyadobacter* sp. MSC1_007**. The conditions are color key are listed. (**A**) 16.F, (**B**) 28.N, (**C**) 39.K, (**D**) 136.osmC, (**E**) 63.S.

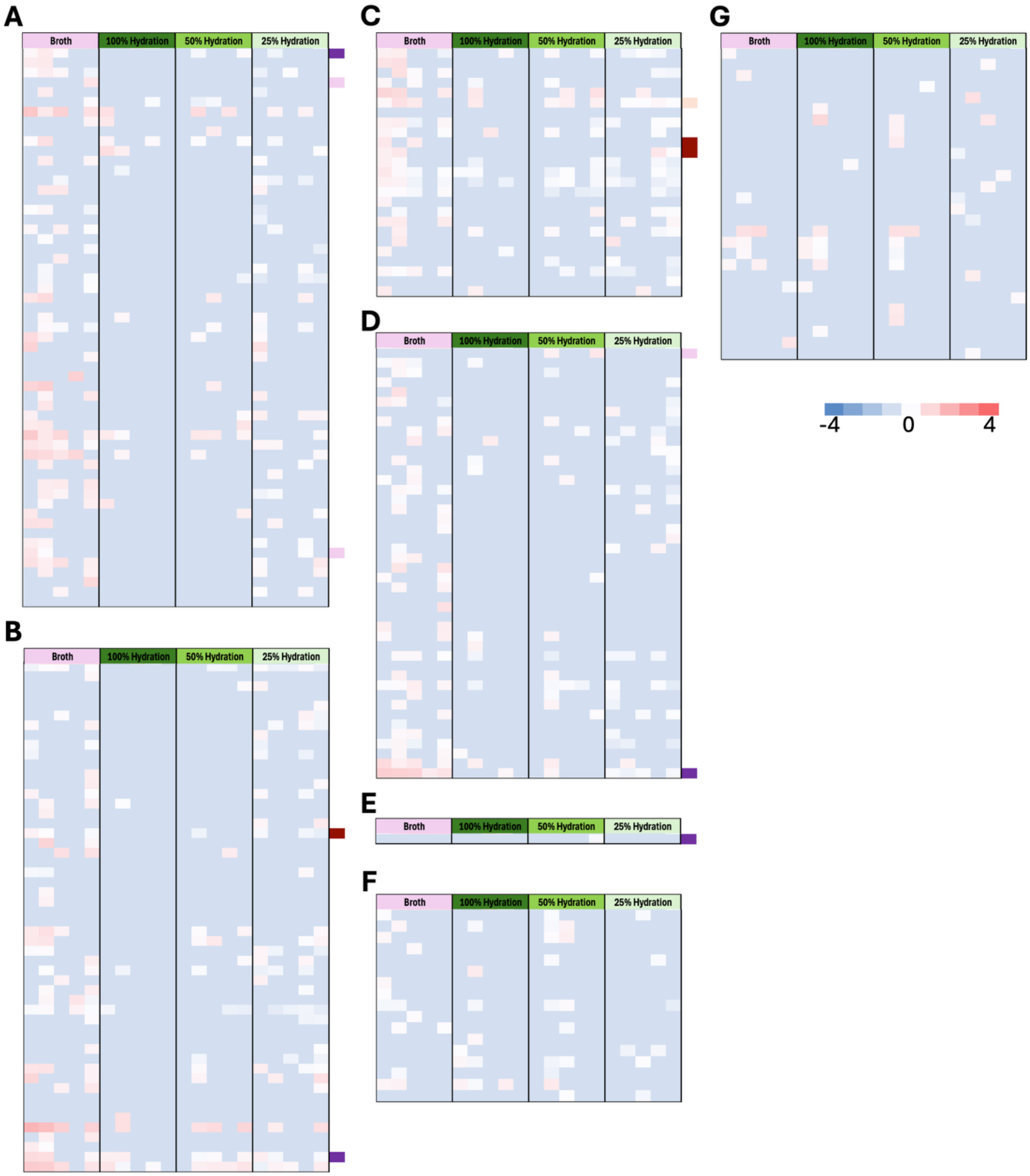


**Supplemental Figure 12. Z-score normalized TPM for the 3 remaining MSC-1 isolate GIs.** The conditions and color keys are listed. (**A-D**) *Neorhizobium tomejilense* MSC1_005 (**A**) 41.T, (**B**) 41.ar14|tig, (**C**) 34.TPP, (**D**) 48.R. (E) *Variovorax beijingensis* MSC1_012 14.yffB. Only a single gene had any TPMs across all conditions and is shown here. (F-G) *Sinorhizobium meliloti* MSC1_014 (**F**) 55.K had 28 predicted genes with TPM and are shown here. (**G**) 47.dusA had 17 genes with TPM shown here.

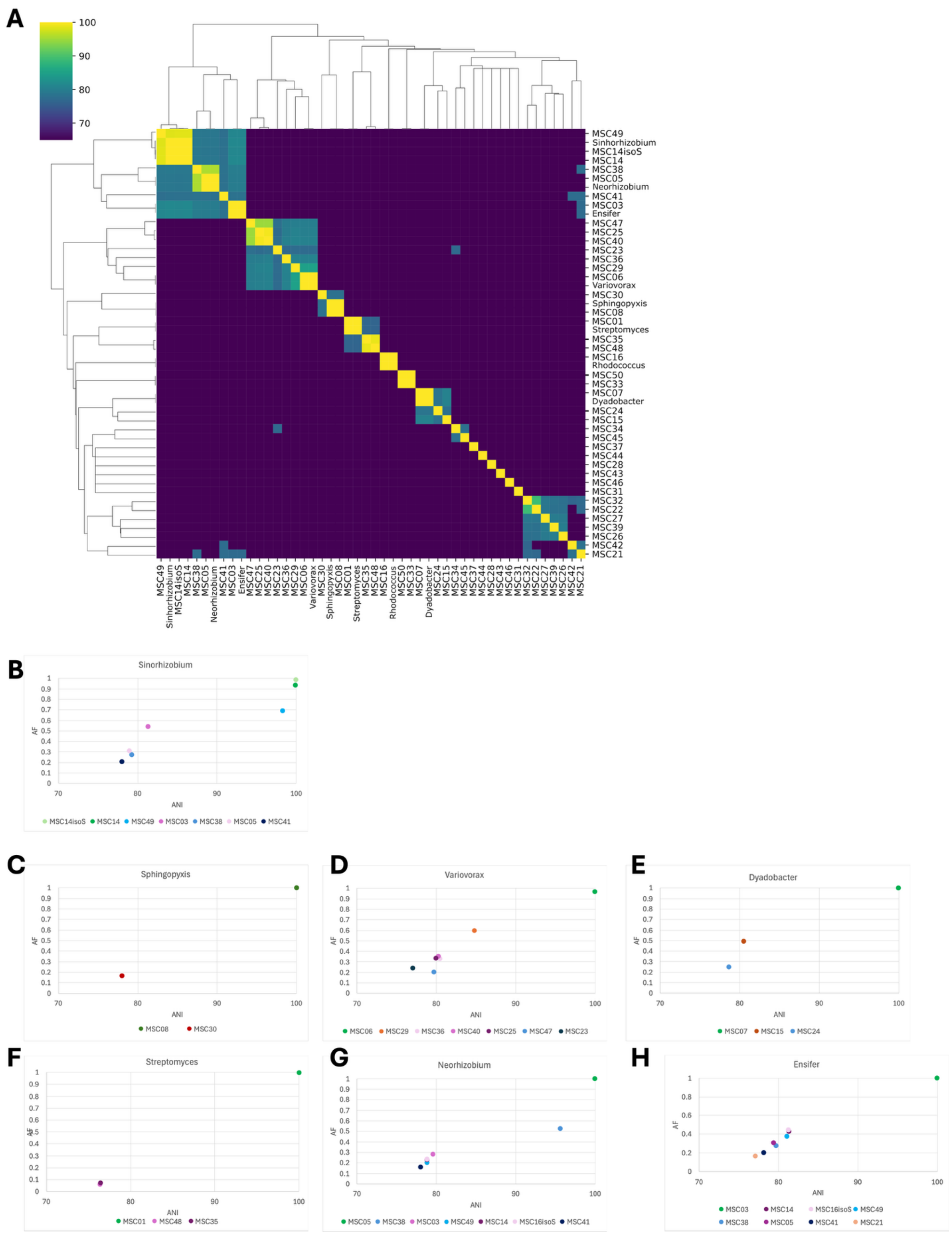


**Supplemental Figure 13. MSC-1 isolate genomes similarity with full MSC-1 community.** (**A**) Hierarchical heatmap of ANI value for MSC-1 genomes. The eight MSC-1 isolates are listed as their genus name. The color key is shown in the upper left. (**B-H**) Scatter plots of ANI and AF values for each MSC-1 isolate compared to any related genomes in the full MSC-1 community.
